# Brassinosteroid receptors reveal conserved features of exon-embedded gene regulation

**DOI:** 10.64898/2026.09.21.753065

**Authors:** Raúl Denia-Mondéjar, Paula Brunot-Garau, Xing Ma, Antonio Serrano-Mislata, María Dolores Gómez, David Latrasse, Moussa Benhamed, Christian S. Hardtke, Noel Blanco-Touriñán

**Affiliations:** Instituto de Biología Molecular y Celular de Plantas, Consejo Superior de Investigaciones Científicas - Universitat Politècnica de València, 46022, Valencia, Spain; Université Paris-Saclay, CNRS, INRAE, Univ Evry, Institute of Plant Sciences Paris-Saclay (IPS2), Orsay 91405, France; Department of Plant Molecular Biology, Biophore, UNIL-Sorge, University of Lausanne, CH-1015 Lausanne, Switzerland

**Author notes:** **Corresponding authors:** Christian S. Hardtke, Noel Blanco-Touriñán. These authors contributed equally to this work (Paula Brunot-Garau, Xing Ma).

## Abstract

Precise quantitative and spatial control of plant gene expression is typically attributed to cis-regulatory elements in promoters and other non-coding sequences. The extent to which exon-encoded regulatory information shapes plant gene expression and how remains unclear. Here we used *BRASSINOSTEROID INSENSITIVE 1 (BRI1)* as a model locus and found that it contains an exonic enhancer (EE), which is essential to establish the spatial *BRI1* expression pattern. Its disruption restricts *BRI1* expression and impairs BRI1-dependent root growth, linking exon-encoded regulatory information and development. The *BRI1* EE acts in concert with its promoter, and related receptor kinase genes likewise harbor regulatory information within their coding sequences. Genome-wide surveys show that candidate EEs are associated with highly connected chromatin interaction networks and transcriptionally active loci. Moreover, species comparisons suggest that common features of exon-encoded regulatory information extend across diverse plant lineages. Thus, exon-encoded regulatory information is a vital component of plant gene regulation.

## Introduction

Precise spatiotemporal control of gene expression chiefly relies on cis-regulatory elements that orchestrate transcriptional programs through transcription factor (TF) recruitment and modulation of chromatin accessibility. Traditionally, these regulatory sequences have been associated with promoters, introns and intergenic regions^[1,2]^, whereas coding sequences are primarily viewed as repositories of polypeptide information. However, evidence from various eukaryotic systems suggests that coding regions can contribute to gene regulation [3-5], revealing an additional layer of genome organization. In plants, it remains poorly understood to what extent and how coding sequences contribute to gene regulation. While introns have long been recognized to enhance gene expression [6-9] and sometimes influence spatial expression [7,10-12], only a few studies have assigned regulatory influence on coding sequences [2,13-15]. However, the recent prediction of thousands of candidate exonic enhancers (EEs) in Arabidopsis (*Arabidopsis thaliana*), based on chromatin accessibility and transcription factor binding sites (TFBSs) [16], implies that exonic DNA sequences may modulate gene expression more frequently than commonly assumed.

We previously uncovered an unexpected expression pattern when *BRI1*, the dominant brassinosteroid receptor in Arabidopsis, was expressed under phloem-specific promoters, since expression expanded beyond the phloem into additional root tissues (Fig. 1a) [13,17]. This phenomenon is not restricted to the root apical meristem, as we observed a similarly expanded pattern in the shoot apical meristem (Fig. 1b). Since neither *BRI1* mRNA nor BRI1 protein appear to move between cells [18,19], and synonymous recoding of the *BRI1* coding sequence abolished the expanded expression pattern [13] (Fig. 1a), this suggests that regulatory information is encoded within the *BRI1* gene body sequence [13,17]. This discovery also supports a model where brassinosteroid signaling predominantly operates cell-autonomously as *BRI1* is ubiquitously expressed [17]. Because *BRI1* is an intron-less gene, the data further exemplify that exonic regulatory information can constitute an integral component of gene regulation, raising several fundamental questions: What is the physiological relevance of regulatory information embedded within the *BRI1* coding sequence? Where is this information located, and what are its features? And more broadly, what are the genomic signatures of exon-embedded regulatory information, and to what extent are they evolutionarily conserved across plant lineages? Here, we address these questions by combining mechanistic dissection of the *BRI1* coding sequence with a recently established catalogue of candidate EEs in Arabidopsis.

**Figure 1.**
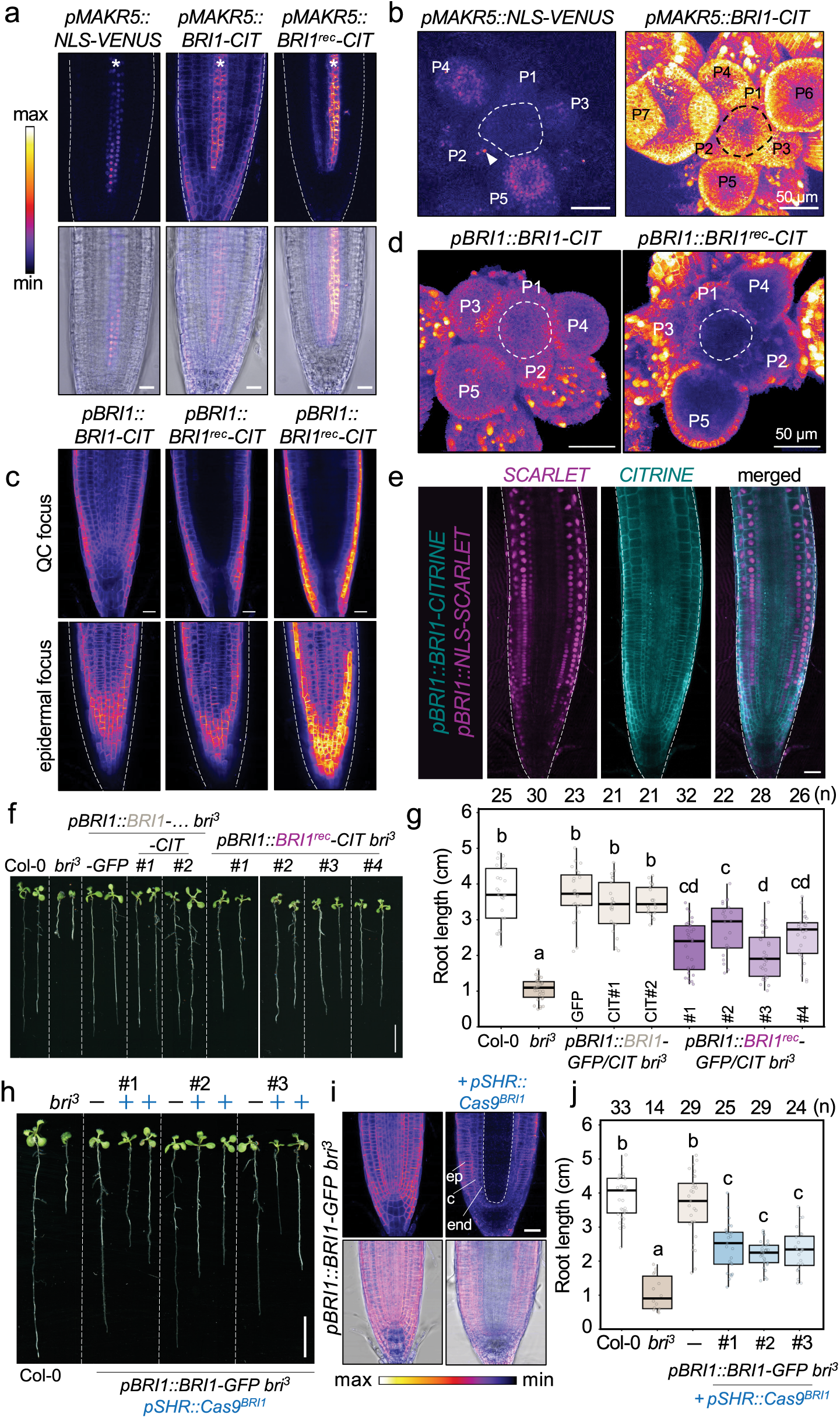
Regulatory information embedded within the coding sequence extends *BRI1* expression to inner root tissues and is required for optimal root growth. **a**, Confocal images of root apices expressing *pMAKR5::NLS–VENUS, pMAKR5::BRI1–CIT*, or the synonymously recoded construct *pMAKR5::BRI1*^*rec*^*–CIT*. Upper panels show fluorescence intensity in false color and lower panels show merged fluorescence and transmitted-light images. Asterisks denote the phloem. Dashed lines outline the root. Scale bars, 20 µm. **b**, Confocal images of shoot apical meristems expressing *pMAKR5::NLS–VENUS* or *pMAKR5::BRI1–CIT*. P1–P7 denote successive floral primordia. The dashed circle indicates the shoot apical meristem. Arrowheads indicate *pMAKR5*-driven NLS–VENUS signal. Scale bars, 50 µm. **c**, Confocal images of root apices expressing *pBRI1::BRI1–CIT* or two independent *pBRI1::BRI1*^*rec*^*–CIT* lines. Upper and lower panels show sections through the quiescent center and epidermis, respectively. The middle *BRI1rec)–CIT* line exhibits fluorescence levels comparable to BRI1–CIT, whereas the right-hand line shows substantially higher fluorescence. Fluorescence is shown in false color. Dashed lines outline the root. Scale bars, 20 µm. **d**, Confocal images of shoot apical meristems expressing *pBRI1::BRI1–CIT* or the higher-expressing *pBRI1::BRI1*^*rec*^*–CIT* line shown in c. P1–P5 denote successive floral primordia. The dashed circle marks the central zone. Scale bars, 50 µm. **e**, Confocal images of root apices co-expressing *pBRI1::NLS–SCARLET* (magenta) and *pBRI1::BRI1–CIT* (cyan). Merged images are shown on the right. Dashed lines outline the root. Scale bars, 20 µm. **f**, Representative 8-day-old seedlings of the indicated genotypes. **g**, Quantification of primary root length of the genotypes shown in f. Each dot represents one seedling. Boxes indicate the interquartile range, center lines indicate medians, and whiskers extend to 1.5 × IQR. Sample sizes (*n*) are indicated above each box. Different letters indicate statistically significant differences between groups (one-way ANOVA followed by Tukey’s HSD test, *P* < 0.05). **h**, Representative 8-day-old seedlings of *pBRI1::BRI1–GFP bri*^*3*^ plants with or without stele-specific *BRI1* editing driven by *pSHR::Cas9BRI1*. Three independent edited lines are shown. **i**, Root apices of *pBRI1::BRI1–GFP bri*^*3*^ plants with or without *pSHR::Cas9BRI1*. Upper panels show fluorescence intensity in false color and lower panels show merged fluorescence and transmitted-light images. Cell layers are indicated (ep, epidermis; c, cortex; en, endodermis). Dashed lines outline the stele. Scale bars, 20 µm. **j**, Quantification of primary root length of the genotypes shown in h. Each dot represents one seedling. Boxes indicate the interquartile range, center lines indicate medians, and whiskers extend to 1.5 × IQR. Sample sizes (*n*) are indicated above each box. Different letters indicate statistically significant differences between groups (one-way ANOVA followed by Tukey’s HSD test, *P* < 0.05).

## Results and Discussion

### Regulatory information embedded within the *BRI1* coding sequence is required for its comprehensive spatial expression

To determine how regulatory information embedded within the *BRI1* coding sequence contributes to its endogenous spatial expression, we first compared expression patterns conferred by the *BRI1* promoter on a nuclear-localized reporter *(pBRI1::NLS-SCARLET)*, a BRI1-CITRINE fusion protein *(pBRI1::BRI1-CIT)*, and a recoded BRI1-CITRINE fusion protein (*pBRI1::BRI1*^*rec*^*-CIT*; a synonymously recoded *BRI1* which preserves the amino acid sequence while c. 25% of the nucleotides are exchanged [13]). Whereas *pBRI1::BRI1-CIT* faithfully recapitulated the reported ubiquitous *BRI1* expression pattern [20] coherent with single cell mRNA sequencing (Fig. S1a) [13], *pBRI1::BRI1*^*rec*^*-CIT* distribution was markedly more restricted: in both root and shoot apical meristems, fluorescence was confined to the outer cell layers (Fig. 1c-d). Within the root apical meristem, BRI1^rec^-CIT expression was predominantly detected in the epidermis and cortex (Fig. 1c). A similarly restricted pattern was observed for *pBRI1::NLS-SCARLET* when co-expressed with either *pBRI1::BRI1-CIT* or *pBRI1::BRI1-GFP*, with NLS-SCARLET fluorescence largely confined to the outer cell layers and weakly extending into the endodermis (Fig. 1e; Fig. S1b,c). This restriction was independent of transgene expression level: even in lines displaying very high overall fluorescence, BRI1^rec^-CIT remained excluded from the inner, vascular tissues, although weak signal was detected in the pericycle (Fig. S1d). Consistently, *pBRI1::NLS-SCARLET* in wild-type seedlings also exhibited fluorescence in the pericycle of strongly expressing lines, but no expression in the vascular tissues (Fig. S1e). These results indicate that the *BRI1* coding sequence is required to recapitulate the full spatial expression pattern of *BRI1*.

We next asked whether *BRI1* expression in inner tissues is functionally important for root growth. Although *pBRI1::BRI1*^*rec*^*-CIT* largely rescued the *bri*^*3*^ root growth defects, quantitative analysis across independent lines revealed that root growth was consistently restored less efficiently than with the *pBRI1::BRI1-GFP/CIT* constructs (Fig. 1f-g). Because complementation efficiency varied across lines, we sought further evidence using a tissue-specific knockout strategy in which CRISPR/Cas9 was expressed from the vascular *SHORT-ROOT (SHR)* promoter to disrupt *BRI1* in the vascular cylinder of the *pBRI1::BRI1-GFP bri*^*3*^ background. In every edited line analyzed, BRI1-GFP fluorescence was lost from the inner tissues but remained detectable in the outer cell layers, and this alteration was consistently accompanied by reduced root growth (Fig. 1h-j).

Altogether, these findings demonstrate that promoter sequences alone are insufficient to recapitulate the *BRI1* expression domain. Instead, regulatory information embedded within the *BRI1* coding sequence provides an additional layer of spatial regulation that extends *BRI1* expression into the inner root tissues, where it is required for optimal root growth. This regulatory architecture may enable *BRI1* expression to be differentially tuned across tissues to maintain appropriate tissue-specific responses [21], with coding-sequence-dependent regulation providing moderate BRI1 levels in the vasculature, where excessive brassinosteroid signaling induces supernumerary cell divisions and radial expansion of the root meristem [22].

### Regulatory information within the *BRI1* coding sequence operates at the transcriptional level

Having established that regulatory information embedded within the coding sequence is required for comprehensive *BRI1* function, we next sought to determine its nature. Several mechanisms could account for our observations. For example, the coding sequence could contain cryptic promoter elements that generate alternative transcripts, influence protein accumulation through sequence-dependent effects on mRNA stability or translation, or instead function as *bona fide* transcriptional cis-regulatory elements.

We first investigated whether the expanded expression pattern could arise from alternative transcription initiation since sequence inspection identified promoter-like motifs, including TATA boxes, that were lost upon synonymous recoding (Fig. S2a). However, neither CAGE-seq (Cap Analysis of Gene Expression and sequencing), monitoring 5’ ends of capped RNA transcripts, nor csRNA-seq (capped small RNA sequencing) revealed evidence of transcription initiation within the *BRI1* gene body (Fig. S2b). Consistently, the *BRI1* coding sequence failed to drive detectable expression when placed downstream of a minimal *35S* promoter, both in transient *Nicotiana benthamiana* leaf assays and in Arabidopsis transformants (Fig. S2d,e). Moreover, this construct failed to rescue the *bri*^*3*^ mutant (Fig. S2c), indicating that the *BRI1* gene body requires promoter context for its regulatory activity.

Next, we considered whether synonymous recoding altered translation efficiency through effects on codon usage, which would provide an alternative explanation for the apparent expression differences [23,24]. However, synonymous recoding did not alter the canonical translational start site, and *BRI1*^*rec*^ produced readily detectable protein when expressed from several tissue-specific promoters across root cell layers (Fig. S3), indicating that the recoded sequence is robustly translated across diverse cellular contexts. Furthermore, *BRI1* expression in inner tissues could reflect inheritance of highly stable *BRI1* mRNA, which may have been abolished by recoding. However, an expanded expression pattern was also observed in the shoot apical meristem when *BRI1* was expressed under the *MEMBRANE-ASSOCIATED KINASE REGULATOR 5* promoter*(pMAKR5)* [25], which is inactive in proliferating shoot meristem cells and only confers expression in developing primordia (Fig. 1b). Collectively, these experiments largely excluded alternative transcription initiation, codon usage-dependent effects on translation, and lineage-dependent inheritance as explanations for the observed discrepancies, rather supporting a transcriptional mechanism.

### An exonic enhancer within the coding sequence confers ubiquitous *BRI1* expression

To map the transcriptional regulatory activity within the *BRI1* gene body, we built on our previous observation that the intracellular region of *BRI1* confers expanded expression when fused to the extracellular domain of BAK1-INTERACTING RECEPTOR-LIKE KINASE 3 (BIR3) (*BIR3*^*ext*^*BRI1*^*int*^) and expressed under the phloem-specific *CLAVATA3/EMBRYO SURROUNDING REGION 45* (*pCLE45)* promoter [13,26]. We independently confirmed this observation using *pMAKR5*, which is specifically active in root meristem phloem poles [25,27] (Fig. 1a). Because this region contains several methylated cytosines [28,29] (Fig. S4a), we tested whether DNA methylation contributes to its activity. Treatment of *pMAKR5::BRI1-CIT* seedlings with the DNA methyltransferase inhibitor 5-azacytidine reduced primary root growth as expected [30] but did not alter the expanded expression pattern (Fig. S4b,c), arguing against DNA methylation as the underlying mechanism.

We therefore scanned the *BRI1* coding sequence for candidate cis-regulatory elements with a focus on enhancers, which can reside within exons [1,16]. Integration of predicted TFBSs with publicly available STARR-seq (Self-Transcribing Active Regulatory Region sequencing) datasets identified a discrete portion of the *BRI1* coding sequence with clustered TFBSs and STARR-seq peaks, which we designated Region 1 (Fig. 2d). The remaining coding sequence was designated Region 2 and encompassed the intracellular domain present in the *BIR3*^*ext*^*BRI1*^*int*^ chimera. Surprisingly, reciprocal BRI1/BRI1^rec^ constructs, in which either Region 1 or Region 2 was synonymously recoded, showed that only recoding Region 1 abolished expression beyond the *pMAKR5* domain (Fig. 2e). To test whether Region 1 functions as a modular cis-regulatory element, we created additional hybrids with the coding sequence of the receptor kinase *BARELY ANY MERISTEM 3 (BAM3)* [26]. Replacement of the BRI1 intracellular domain with the corresponding BAM3 region still conferred broad expression throughout the root under *pBRI1*, whereas the reciprocal construct remained restricted to the outer tissues (Fig. S5). Thus, the regulatory activity associated with Region 1 is transferable to a heterologous coding-sequence context and functions independent of the surrounding *BRI1* coding sequence, consistent with a modular exonic enhancer.

**Figure 2.**
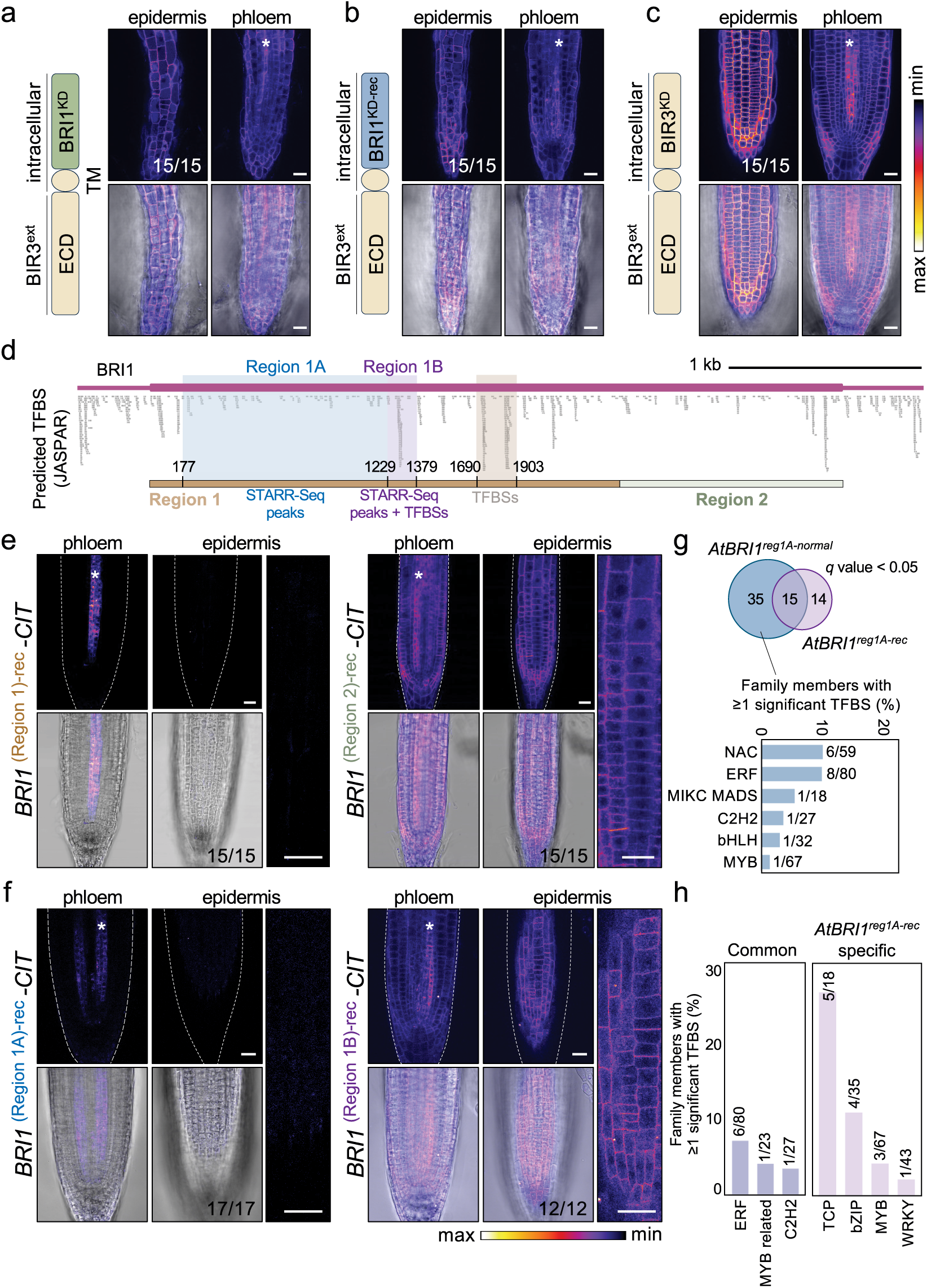
Mapping of a coding sequence interval required for the expanded expression pattern of BRI1 under tissue-specific promoters. **a**, Confocal images of root apices expressing a *pMAKR5::BIR3*^*EXT*^*–BRI1*^*INT*^*–CITRINE* chimeric receptor. The extracellular and transmembrane domains (ECD and TM) derive from BIR3, whereas the intracellular kinase domain (KD) derives from BRI1. Images were acquired in the epidermal and phloem focal planes. Upper panels show fluorescence intensity in false color and lower panels show merged fluorescence and transmitted-light images. The number of independent transformants displaying the representative expression pattern is indicated. Asterisks denote the phloem. Scale bars, 20 µm. **b**, Confocal images of root apices expressing the synonymously recoded *pMAKR5::BIR3*^*EXT*^*–BRI1*^*INTrec*^*–CITRINE* chimera. Images are displayed as in a. **c**, Confocal images of root apices expressing *pMAKR5::BIR3–CITRINE*. Images are displayed as in a. **d**, Predicted transcription factor binding sites (TFBSs; JASPAR) across the *BRI1* coding sequence together with publicly available STARR-seq peaks. Two candidate regions (Regions 1 and 2) and the corresponding subregions (Regions 1A and 1B within Region 1) used to generate synonymously recoded constructs are indicated. **e**, Confocal images of root apices expressing *pMAKR5::BRI1(Region 1)*^*rec*^*–CIT* or *pMAKR5::BRI1(Region 2)*^*rec*^*–CIT*. Images were acquired in the phloem and epidermal focal planes. Upper panels show fluorescence intensity in false color and lower panels show merged fluorescence and transmitted-light images. Right panels show higher-magnification views of the epidermis. Asterisks denote the phloem. Dashed lines outline the root. The number of independent transformants displaying the representative expression pattern is indicated. Scale bars, 20 µm. **f**, Confocal images of root apices expressing *pMAKR5::BRI1(Region 1A)*^*rec*^*–CIT* or *pMAKR5::BRI1(Region 1B)*^*rec*^*–CIT*. Images are displayed as in e. **g**, Comparison of predicted TF representation between *BRI1* Region 1A native and synonymously recoded sequences. The Venn diagram shows shared and sequence-specific TFs with at least one significant TFBS (q < 0.05). The bar plot shows TF family representation among TFs specific to the native sequence, calculated as the percentage of TFs within each family represented by at least one significant TFBS. **h**, TF family representation among TFBSs shared between the native and recoded Region 1A sequences (Common) or specific to the recoded sequence. TF family representation was calculated as the percentage of TFs within each family represented by at least one significant TFBS (q < 0.05).

To further refine the EE location, we generated additional BRI1/BRI1^rec^ hybrids in which either the STARR-seq peak-containing subregion (Region 1A) or the adjacent STARR-seq peak- and TFBS-rich subregion (Region 1B) was synonymously recoded while the remainder of Region 1 retained the native sequence (Fig. 2d). These experiments mapped the EE to Region 1A as the subregion responsible for the expanded *BRI1* expression pattern (Fig. 2f). Region 1A contains predicted binding sites for numerous TF families, many of which were altered by synonymous recoding (Fig. 2g-h). Whether the regulatory activity of Region 1A depends on combinatorial organization of multiple TFBSs or instead on a few functionally dominant motifs remains to be determined.

Having mapped the *BRI1* EE to Region 1A, outside the intracellular coding region present in the *BIR3*^*ext*^*BRI1*^*int*^ chimera, we revisited the basis of the expanded expression observed with this construct. We reasoned that this pattern might instead reflect regulatory activity within the *BIR3* coding sequence. Synonymous recoding of the intracellular BRI1 region in the *BIR3*^*ext*^*BRI1*^*int*^ hybrid did not abolish the expanded pattern (Fig. 2b), and expression of *BIR3* itself under *pMAKR5* also expanded beyond the phloem (Fig. 2c). Thus, *BIR3* apparently contains coding sequence-embedded regulatory information of its own, raising the possibility that such regulatory activity may be common across Arabidopsis genes.

### Exonic enhancers are widespread across Arabidopsis genes and define a distinct regulatory architecture

To assess the prevalence of coding sequence-embedded regulatory activity genome wide, we leveraged a recent study that identified thousands of candidate exonic enhancers (cEEs) in Arabidopsis based on chromatin accessibility, enhancer-associated chromatin signatures, and dense clusters of predicted TFBSs [16]. However, beyond the features used for their initial prediction, little is known about their shared molecular and genomic properties. We therefore systematically analyzed the Arabidopsis catalogue to identify common characteristics of cEEs and determine whether the *BRI1* EE conforms to these broader patterns. The catalogue comprises 7,138 cEEs distributed across 5,857 genes [16], with most genes containing a single cEE and a smaller subset harboring multiple candidate elements (Fig. 3a,b). Gene Ontology analysis revealed that cEE-containing genes are enriched for biological processes associated with environmental responses and development (Fig. 3c). Although present in diverse genes, cEEs showed preferential associations with specific TF families. Several TF families, including ERF, bZIP, WRKY and NAC, were particularly well represented among cEEs (Fig. 3d, Fig. S6). Consistent with this distinctive TFBS composition, synonymous *BRI1* recoding depleted binding sites of cEE-enriched TF families, while the largest gain corresponded to TCP binding sites, which were less enriched among cEEs than other TF families (Fig. 2g,h; Fig. 3d). Because each cEE corresponds to an annotated exon rather than a precisely mapped regulatory sequence, its position within the coding sequence can only be approximated in the absence of functional mapping. We therefore assigned each cEE the midpoint of its annotated exon and examined the positional distribution separately for genes in which the cEE-containing exon occupied different fractions of the gene. Across all size classes, cEE-containing exons were distributed throughout genes but consistently showed a relative enrichment towards the 3’ portion of genes (Fig. 3e,f).

**Figure 3.**
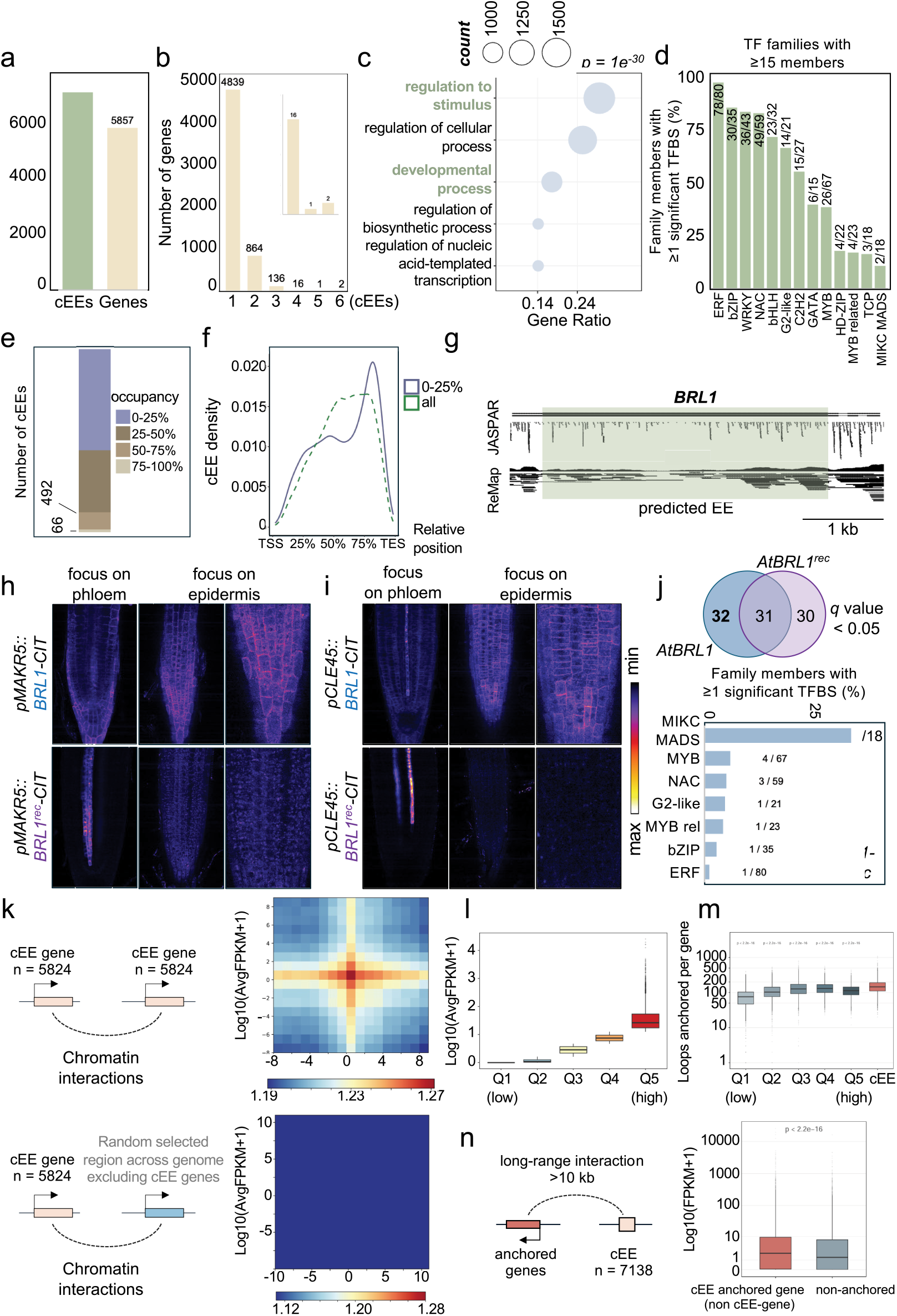
Genome-wide properties of candidate exonic enhancers in Arabidopsis. **a**, Number of candidate exonic enhancers (cEEs) and genes containing cEEs in the Arabidopsis genome. **b**, Distribution of cEEs per gene. **c**, Gene Ontology enrichment analysis of genes containing cEEs. The most significantly enriched biological processes are shown. Bubble size represents the number of genes associated with each GO term, and the x-axis indicates the gene ratio. **d**, TF family representation in cEEs, based on motif scanning using PlantTFDB. TF families comprising at least 15 members are shown; families with fewer than 15 members are shown in Extended Data Fig. 6. Numbers indicate the number of TFs in each family for which at least one significant TFBS was predicted relative to the total number of TFs in that family; bar height represents the corresponding percentage. **e**, Distribution of the fraction of gene length occupied by cEEs. cEEs were grouped according to the fraction of gene length they occupy. **f**, Metagene analysis showing the positional distribution of cEEs relative to gene length. Curves show all cEEs or those occupying 0–25% of gene length. **g**, Predicted TFBSs across the *BRL1* coding sequence. The predicted EE is indicated. **h**, Confocal images of root apices expressing *pMAKR5::BRL1–CIT* or the synonymously recoded *pMAKR5::BRL1*^*rec*^*–CIT*. Images were acquired in the phloem and epidermal focal planes. Right panels show higher-magnification views of the epidermis. The number of independent transformants displaying the representative expression pattern is indicated. Asterisks denote the phloem. Dashed lines outline the root. Scale bars, 20 µm. **i**, Confocal images of root apices expressing *pCLE45::BRL1–CIT* or *pCLE45::BRL1*^*rec*^*–CIT*. Images are displayed as in h. **j**, Comparison of predicted TF representation between the native and synonymously recoded *BRL1* sequences. The Venn diagram shows shared and sequence-specific TFs with at least one significant TFBS (q < 0.05). The bar plot shows TF family representation among TFs specific to the native sequence, calculated as the percentage of TFs within each family represented by at least one significant TFBS. **k**, Schematic of the chromatin interaction analysis workflow and aggregate Micro-C contact maps for pairs of cEE-containing genes and pairs comprising a cEE-containing gene and a randomly selected non-cEE control gene. **l**, Distribution of gene expression levels used to define the five expression quintiles (Q1–Q5) for subsequent chromatin interaction analyses. **m**, Number of chromatin interactions anchored at cEE-containing genes (EE) and genes grouped by expression quintile (Q1–Q5). Statistical significance was assessed using Wilcoxon rank-sum tests comparing cEE-containing genes with each expression quintile. **n**, Expression levels of non-cEE genes anchored to cEE-containing genes through long-range (>10 kb) chromatin interactions (cEE-anchored) or not anchored to cEE-containing genes (non-cEE-anchored). Only chromatin loops with Z-score > 1000 and FDR < 0.05 were considered. Statistical significance was assessed using a Wilcoxon rank-sum test. Numbers indicate the number of genes in each group.

Interestingly, the two other brassinosteroid receptors, *BRI1-LIKE 1 (BRL1)* and *BRL3* also appeared in the cEE catalogue (Fig. 3g, Fig. S7a, Supplemental Table 4). Expression of both *BRL1* and *BRL3* fusion proteins driven by the phloem-specific *pMAKR5* or *pCLE45* promoters extended into surrounding tissues, indicating that coding sequence-embedded regulatory information indeed contributes to their spatial expression (Fig. 3h,i and Fig. S7b). We focused on *BRL1* to test whether this depends on the cEE. Synonymous recoding of *BRL1* indeed abolished expression outside the phloem while preserving expression within the phloem, copying the effect observed for *BRI1*^*rec*^ (Fig. 3h,i). Consistent with this functional disruption, synonymous recoding extensively remodeled the predicted TF binding landscape of the *BRL1* cEE (Fig. 3j). Together, these findings indicate that coding sequence-dependent spatial regulation extends beyond *BRI1* and *BIR3*, and establish *BRL1* and *BRL3* as additional examples for functional EEs. Notably, the endogenous *BRL1* and *BRL3* expression domains are spatially more restricted than *BRI1* [13,31,32], consistent with the idea that EEs cooperate with promoter-proximal regulatory information to shape final expression patterns.

### Exonic enhancers reside within highly connected chromatin interaction networks linking transcriptionally active loci

Beyond the examples above, where coding sequence-embedded enhancers expand spatial expression, the full functional repertoire of cEEs may also include fine-tuning transcriptional output or regulating neighboring genes in *cis* through long-range regulatory interactions (Fig. S8). To begin exploring this broader regulatory potential, we asked whether cEEs are embedded within chromatin interaction networks compatible with regulatory functions beyond their resident genes. Analysis of publicly available high-depth Micro-C datasets from Arabidopsis seedlings revealed extensive chromatin interactions associated with cEE-containing loci, including contacts spanning neighboring genes (Fig. S9a). Consistently, aggregate contact analysis of cEE-containing genes (n=5,284) showed a pronounced enrichment in contact frequency compared with randomly selected genomic regions (Fig. 3k), indicating that cEE-associated loci are not randomly organized within the three-dimensional genome.

Because chromatin connectivity is often associated with transcriptional activity, we asked whether this enrichment simply reflects gene expression. We stratified Arabidopsis genes into five expression quintiles based on RNA-seq data, and 1,000 genes were randomly sampled from each quintile for comparison (Fig. 3l). cEE-containing genes anchored significantly more chromatin loops than genes in any expression quintile (Fig. 3m; *P*<2.2×10^-16^; Fig. S9b), indicating that their enhanced connectivity cannot be explained solely by transcriptional activity. Using high-confidence long-range interactions (>10kb), we identified 15,427 genes connected to cEE-containing loci. These genes showed significantly higher expression than non-associated genes (median 2.26 versus 1.42 FPKM; *P*<2.2×10^-16^) (Fig. 3n), indicating that cEE-associated chromatin interactions preferentially connect transcriptionally active genes. Thus, while EEs can operate locally to regulate their resident genes, their chromatin interactions raise the possibility that they also contribute to regulatory programs of neighboring, even distant loci.

### Exon-embedded regulatory features extend across divergent plant lineages

Having uncovered exon-embedded regulatory information in *BRI1* and defined broader features associated with Arabidopsis cEEs, we investigated whether similar regulatory properties are conserved across plant evolution. We initially focused on a *BRI1* orthologue from the distantly related Brassicaceae *Cochlearia groenlandica (CgBRI1)* [33], which indeed retained the ability to direct expression beyond promoter-defined domains in Arabidopsis. Regardless of whether *CgBRI1* was expressed from *pMAKR5, pBRI1*, or the epidermis-specific *ARABIDOPSIS THALIANA MERISTEM LAYER 1 (ATML1)* promoter, its expression extended beyond the corresponding activity domains (Fig. 4a,b), replicating the behavior of Arabidopsis *BRI1*. Thus, coding sequence-dependent regulation is a conserved property of this *BRI1* orthologue.

**Figure 4.**
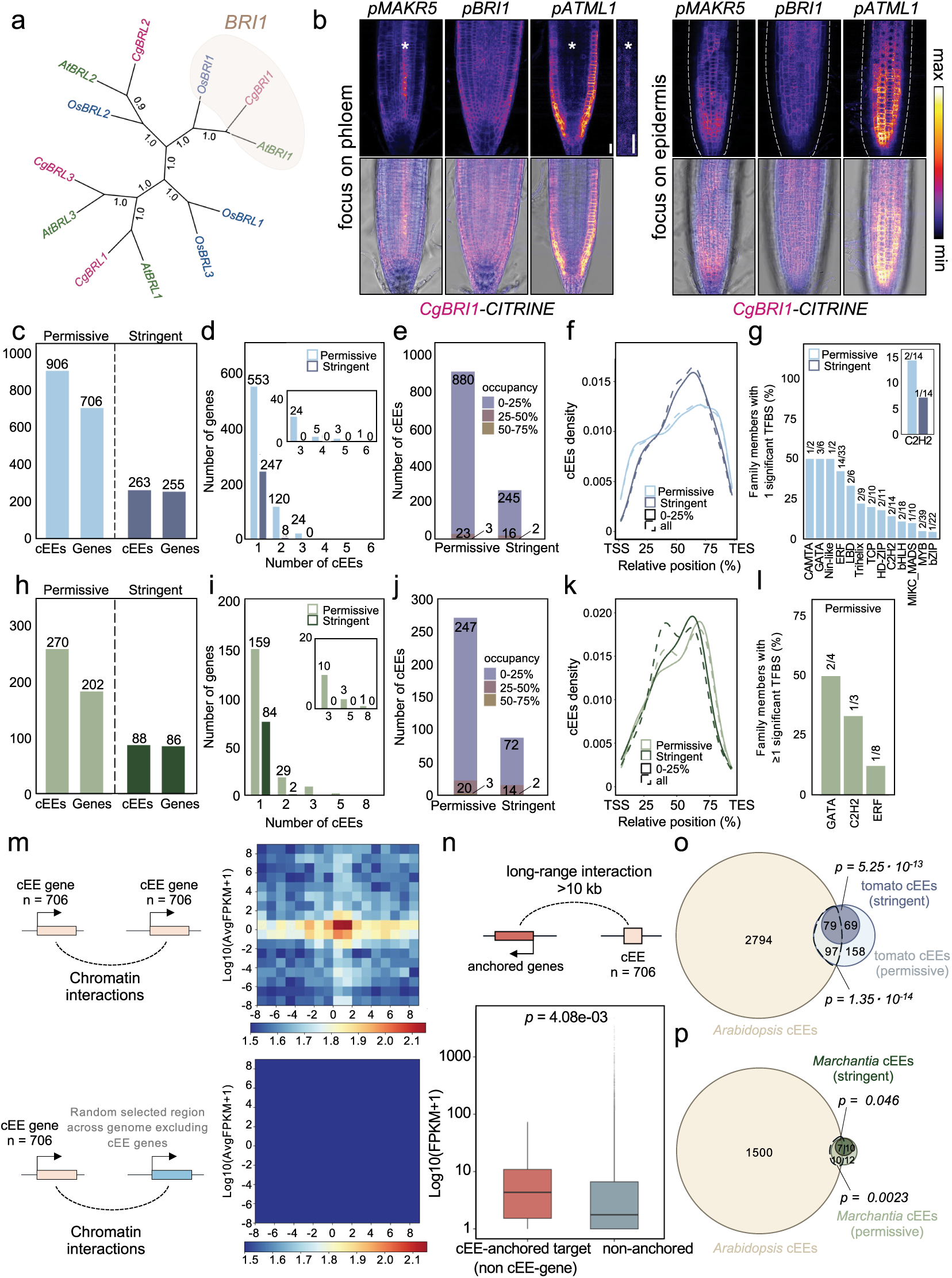
Evolutionary conservation of exon-embedded regulatory features in plants. **a**, Maximum-likelihood phylogeny of *BRI1/BRL* genes from Arabidopsis, *Cochlearia* and rice. The *Cochlearia* BRI1 orthologue (CgBRI1) used for functional analyses is highlighted. **b**, Confocal images of Arabidopsis roots expressing CgBRI1–CITRINE under the control of the phloem-specific *pMAKR5*, native *pBRI1* or epidermis-specific *pATML1* promoters. Left panels show optical sections through the phloem and right panels through the epidermis. Fluorescence intensity is shown in false color (upper) and merged with transmitted-light images (lower). Asterisks denote the phloem. Dashed lines outline the root. Scale bars, 20 µm. c–g, Genome-wide identification and characterization of candidate exonic enhancers (cEEs) in tomato (*Solanum lycopersicum*) using permissive and stringent identification criteria. **c**, Number of identified cEEs and cEE-associated genes. **d**, Distribution of the number of cEEs per gene. **e**, Distribution of the fraction of gene length occupied by cEEs. f, Relative positional distribution of cEEs along genes. **g**, Representation of transcription factor families among cEEs. TF family representation was calculated as the percentage of TFs within each family represented by at least one significant TFBS (*q* < 0.05). h–l, Genome-wide identification and characterization of cEEs in *Marchantia polymorpha* using permissive and stringent identification criteria. **h**, Number of identified cEEs and cEE-associated genes. **i**, Distribution of the number of cEEs per gene. **j**, Distribution of the fraction of gene length occupied by cEEs. **k**, Relative positional distribution of cEEs along genes. **l**, Representation of transcription factor families among cEEs. TF family representation was calculated as the percentage of TFs within each family represented by at least one significant TFBS (*q* < 0.05). **m**, Schematic of the chromatin interaction analysis workflow and aggregate micro-C contact maps for pairs of cEE-containing genes and pairs comprising a cEE-containing gene and a randomly selected non-cEE control region in tomato. **n**, Expression levels of non-cEE genes anchored or not anchored to cEE-containing genes through long-range (>10 kb) chromatin interactions in tomato. Only chromatin loops with Z-score > 1000 and FDR < 0.05 were considered. Statistical significance was assessed using a Wilcoxon rank-sum test. **o**, Expression levels of non-cEE genes anchored or not anchored to cEE-containing genes through long-range (>10 kb) chromatin interactions in tomato. p–q, Overlap between orthogroups containing cEE-associated genes in *Arabidopsis thaliana* and *Solanum lycopersicum* (p) or *Marchantia polymorpha* (q) using permissive and stringent cEE identification criteria. *p* values indicate enrichment above random expectation.

We next expanded our analysis from individual *BRI1* orthologues to the genome-wide characterization of cEEs in tomato *(Solanum lycopersicum)* and the liverwort Marchantia *(Marchantia polymorpha)*. Because the comprehensive regulatory datasets available for Arabidopsis are still lacking in these species, we adapted the cEE identification strategy to the available genomic resources, using chromatin accessibility as a proxy for regulatory potential, although this approach is expected to recover fewer cEEs. Specifically, we identified exons located more than 100 bp from annotated transcription start and termination sites and overlapping ATAC-seq (Assay for Transposase-Accessible Chromatin using sequencing) peaks (permissive), and also defined a more stringent subset by requiring the ATAC-seq peak summit to fall within the exon (stringent). The permissive criterion identified 906 cEEs distributed across 706 tomato genes and 270 cEEs distributed across 202 Marchantia genes, whereas the stringent criterion retained 263 cEEs across 255 tomato genes and 88 cEEs across 86 Marchantia genes (Fig. 4c,h). Tomato cEEs showed signals associated with active chromatin, including H3K4me3 and H3K9ac, while repressive histone marks, such as H3K27me3 and H3K9me2, showed little signal around their centers (Fig. S10a). Interestingly, these active chromatin signatures were less pronounced than at the transcription start sites (TSSs) of their host genes, particularly for H3K4me3 (Fig. S10a,b). As in Arabidopsis, cEE-containing genes in tomato and Marchantia typically harbored a single cEE, and cEEs occupied a relatively small fraction of the coding sequence (Fig. 4d,e,i,j). Moreover, cEEs displayed broadly similar positional distributions across species, with a tendency to accumulate in the second half of coding sequences (Fig. 4f,k). Importantly, these genomic features remained largely unchanged under the stringent criterion. TFBS representation also showed similarities across species, with all TF families identified in tomato and Marchantia also represented in Arabidopsis cEEs. This pattern was maintained under the stringent criterion, despite the loss of significant TFBS enrichment in both tomato and Marchantia (Fig. 3d, Fig. 4 g,l; Fig. S6).

Aggregate contact analysis in tomato revealed increased chromatin interactions around cEE-containing loci compared with randomly selected genomic regions, mirroring the pattern observed in Arabidopsis (Fig. 4m). To assess the contribution of transcriptional activity to this increased connectivity, we stratified genes into expression quintiles and quantified both the number of loops anchored by each gene and the maximum Z-score of these interactions. Both measures generally increased with gene expression, with cEE-containing genes displaying connectivity profiles comparable to those of highly expressed genes (Fig. S10c-e). Thus, transcriptional activity cannot be excluded as a contributor to the increased chromatin connectivity associated with tomato cEEs. Notably, non-cEE genes connected to cEE-containing loci were more highly expressed than non-connected genes, an association that remained significant when the analysis was restricted to high-confidence long-range interactions (>10 kb; *P*=4.41×10^−3^) (Fig. 4n). Together, these analyses reveal broad similarities in the genomic and chromatin features associated with cEEs across species. Yet, it remained unclear whether the association of cEEs with particular gene families is conserved across species. To address this, we examined whether cEE-associated genes were enriched in shared orthogroups. Using the permissive criterion, we identified 176 orthogroups containing cEE-associated genes in both Arabidopsis and tomato, significantly more than expected by chance (*P*=1.35×10^−14^) (Fig. 4o). Notably, 17 orthogroups contained cEE-associated genes in both Arabidopsis and Marchantia, again exceeding random expectation (*P*=0.0023) (Fig. 4p). This enrichment was retained under the stringent criterion in both comparisons, with 79 Arabidopsis–tomato and seven Arabidopsis– Marchantia shared orthogroups (*P*=5.35×10^−13^ and *P*=0.046, respectively) (Fig. 4o,p). The enrichment of shared cEE-associated orthogroups above random expectation suggests that the propensity to harbor cEEs may be evolutionarily conserved in a subset of gene families. Together with the conserved regulatory behavior of *BRI1*, these findings support shared features of exon-embedded regulation across plant lineages.

Our functional and genome-wide analyses raise the possibility that coding sequence-embedded regulatory information may be more widespread than commonly assumed. A few studies have reported spatially expanded expression patterns of translational reporters when compared to transcriptional reporters [14,15,34] and sometimes excluded mobile mRNA or protein as an explanation [14,15]. A systematic survey of transgenic reporters may reveal the frequency of such discrepancies and their association with cEEs. Our cross-species analyses further suggest that some of these features extend across divergent plant lineages. Yet, the regulatory scope of coding sequences may be even broader than explored here. For instance, whether coding sequences also harbor repressive regulatory elements remains an outstanding question. To our knowledge, the only functionally characterized examples of repressive cis-regulatory elements embedded within plant coding sequences are the BSM1 elements, first identified in the *Gynandropsis gynandra NAD-dependent MALIC ENZYME1* coding sequence [35]. These elements establish bundle sheath-specific expression by repressing transcription in neighboring mesophyll cells. The increasing availability of genome-wide functional datasets provides an opportunity to uncover such repressive regulatory elements. Such efforts could define the full repertoire of regulatory information embedded within coding sequences and clarify how this additional layer of gene regulation contributes to plant development and evolution beyond brassinosteroid signaling.

## Methods

### PLANT material and growth conditions

All lines used in this work were in the Arabidopsis wild-type accession Columbia-0 (Col-0) background. The *bri1-116 brl1 brl3* triple mutant (*bri*^*3*^) and the *pMAKR5::NLS-VENUS, pMAKR5::BRI1-CIT bri*^*3*^, *pMAKR5::BRI1*^*rec*^*-CIT bri-116 (+/-) brl1 brl3, pBRI1::BRI1-CIT bri*^*3*^, *pBRI1:BRI1-GFP bri*^*3*^, *pBRI1::BRI1*^*rec*^*-CIT bri*^*3*^, *pCLE45::BRL1-CIT bri*^*3*^, *pCLE45::BRL3-CIT bri*^*3*^, *pATML1::BRI1*^*rec*^*-CIT bri-116 (+/-) brl1 brl3*, and *pSHR::BRI1*^*rec*^*-CIT bri-116 (+/-) brl1 brl3* transgenic lines were described previously [13,22,36]. For tissue culture, seeds were surface sterilized, stratified at 4 °C for 2 days and grown vertically on half-strength Murashige and Skoog (½ MS) medium supplemented with 1% agar and 0.3% sucrose (pH 5.7), at 22 °C under continuous white LED light (∼120 μE). For 5-azacytidine treatments, seeds were germinated and seedlings were grown for 5 days on the same medium supplemented with 10 or 50 μM 5-azacytidine or the corresponding DMSO control. For shoot apical meristem experiments, plants were grown on soil (sphagnum:perlite:vermiculite, 2:1:1) at 21 °C under long-day conditions (16 h light/8 h dark) with cool-white, fluorescent light (∼150 μE m−2 s−1); seeds were stratified on soil for 3 days at 4 °C in darkness before germination. For propagation and seed production, seedlings germinated on MS plates as described above were transferred to soil and grown under the same long-day conditions.

### Construct generation and plant transformation

Full-length *BRI1* coding sequences carrying synonymous recoding of Region 1, Region 1A, Region 1B or Region 2, together with *BIR3*^*EXT*^*– BRI1*^*INT-rec*^, *BRL1*^*rec*^ and *BZR1*, were synthesized and cloned directly into pTWIST entry vectors by Twist Bioscience. *BIR3* and *BRI1*^*EXT*^*–BAM3*^*INT*^ were synthesized separately and introduced into pDONR221 by BP recombination (Invitrogen). *CgBRI1* and *BREVIS RADIX-LIKE 4 (BRXL4)* were PCR-amplified using the primers listed in Supplementary Table 2 and introduced into pDONR221 by BP recombination. Entry clones containing *BRI1, BRI1*^*rec*^, *BRL1, BRL3, BAM3*^*EXT*^*–BRI1*^*INT*^ and *BIR3*^*EXT*^*– BRI1*^*INT*^ were described previously [13,37]. Coding sequences were assembled without their native stop codons for in-frame C-terminal fusion to CITRINE.

Gateway P4-P1R entry clones containing the *BRI1, MAKR5, ATML1, CORTEX 2 (CO2), PINFORMED 2 (PIN2), CASPARIAN STRIP 1 (CASP1), SHR* and *CLE45* promoters were generated and used previously [13,25,26,38-41]. The minimal *35S* promoter *(35Smin)* was PCR-amplified using primers [35Smin-F] and [35Smin-R] (Supplementary Table 2) and introduced into the Gateway P4-P1R entry vector by BP recombination. Promoter and coding sequence entry clones were combined with the appropriate fluorescent reporter modules by Multisite Gateway LR recombination (Thermo Fisher Scientific). The *pBRI1::NLS-SCARLET* construct was assembled in the two-fragment destination vector pFR7m24GW, whereas promoter–CDS–reporter combinations were assembled in pFR7m34GW (FAST RED) or pFG7m34GW (FAST GREEN). Final constructs were introduced into *Agrobacterium tumefaciens* strain GV3101 carrying the pMP90 helper plasmid, and Arabidopsis plants were transformed by floral dip.

### Tissue-specific CRISPR targeting of BRI1

Tissue-specific CRISPR/Cas9 targeting of *BRI1* was performed using the previously described *pSHR::Cas9*^*BRI1*^ construct [13]. Independent *pBRI1::BRI1-GFP bri*^*3*^ *pSHR::Cas9*^*BRI1*^ lines were analyzed by confocal microscopy and root-growth assays.

### Root growth analysis

Eight-day-old seedlings were imaged on growth plates using a high-resolution flatbed scanner, and primary root length was measured from the hypocotyl–root junction to the root tip using Fiji software. For representative images, seedlings were transferred to MS plates containing India ink and photographed. Outliers were identified separately within each genotype using the 1.5 × interquartile range (IQR) criterion and excluded before analysis. Genotypes were compared by one-way ANOVA followed by Tukey’s HSD test (*P* < 0.05). Boxplots show medians and IQRs, with whiskers extending to 1.5 × IQR and individual observations shown as points. Sample sizes (*n*) indicate observations included after outlier filtering. Statistical analyses and plotting were performed in R using ggplot2. Experiments were independently repeated at least twice with comparable results.

### Confocal microscopy and image processing

Shoot apices were excised from inflorescences bearing approximately three open flowers, and floral buds were carefully removed under a stereomicroscope using a fine needle to expose the shoot apical meristem. Z-stacks of the shoot apices were acquired on a Leica Stellaris 8 FALCON confocal microscope with a 0.5-μm step size. Longitudinal optical sections of primary roots were acquired using Leica Stellaris 5 and Zeiss LSM 780 confocal microscopes. Images shown in Figs. 1a, 2a–c, 2f, Fig. S1b and Fig. S2d,e were acquired using the Zeiss LSM 780; all other confocal images were acquired using Leica Stellaris microscope.

For fluorescence imaging on the Leica Stellaris microscopes, GFP was excited at 488 nm, whereas CITRINE and NLS-VENUS were excited at 514 nm; emission was collected at 493–565 nm. NLS-SCARLET was excited at 561 nm and emission was collected at 566–734 nm. On the Zeiss LSM 780, CITRINE and NLS-VENUS were excited at 514 nm using an argon laser, and emission was collected at 518–616 nm. Laser power and detector gain were adjusted to maximize fluorescence signal while avoiding saturation and were kept constant across samples compared within the same experiment. Images were processed and analyzed using Fiji (ImageJ) software [42]. Z-stack projections and adjustments to brightness and contrast were performed in Fiji. Fluorescence intensity profiles in Fig. S1c were obtained in Fiji from a rectangular ROI spanning the root width. The same ROI was applied to the GFP and NLS-SCARLET channels, and mean fluorescence intensity along the ROI was obtained using the Plot Profile function. Profiles were exported to R, independently min–max normalized (0–1) for each channel and plotted as a function of distance across the root.

### Minimal-promoter assays

The *35Smin::BRI1-CITRINE* construct was analyzed in more than 20 independent T1 Arabidopsis transformants in each of the Col-0 and *bri*^*3*^ backgrounds. The presence of the transgene in individual T1 seedlings was confirmed by PCR genotyping. CITRINE fluorescence was assessed in Col-0, whereas complementation in the *bri*^*3*^ background was evaluated based on restoration of root growth and rescue of the characteristic cabbage-like phenotype. For transient expression assays, leaves of one-month-old *Nicotiana benthamiana* plants grown at 25°C under a 16-h light/8-h dark photoperiod were infiltrated with *Agrobacterium tumefaciens* GV3101 carrying *35Smin::BRI1-CITRINE* together with the p19 silencing suppressor. A *35S::GFP* construct was infiltrated in parallel as an infiltration control. Bacterial suspensions carrying *35Smin::BRI1-CITRINE, 35Smin::BRI1*^*rec*^*-CITRINE* or *35S::GFP* and p19 were adjusted to final OD600 values of 0.2 and 0.05, respectively. Reporter expression was analyzed by confocal microscopy three days after infiltration, with more than 10 leaf discs examined per construct.

### Analysis of CAGE-seq, csRNA-seq and ATAC-seq datasets

For Arabidopsis, normalized genome-wide coverage tracks in BigWig format were obtained from previously published CAGE-seq, csRNA-seq and ATAC-seq datasets. Processed CAGE-seq data [43] were downloaded from the Plant Epigenetics portal, whereas csRNA-seq data from seedlings collected after 57 h in the light and ATAC-seq data were obtained from the NCBI Gene Expression Omnibus (GEO; accessions GSE250331 and GSE123263, respectively) [44,45]. Genomic signal tracks were visualized together with the TAIR10 reference genome annotation around genes of interest using Integrative Genomics Viewer (IGV) v2.18.4 [46].

### ATAC-seq data processing and peak calling

For Marchantia, raw paired-end ATAC-seq reads were obtained from the Sequence Read Archive (SRA; accession SRR10879463). Read quality was assessed using FastQC (https://www.bioinformatics.babraham.ac.uk/projects/fastqc/), and adapter trimming and quality filtering were performed with Trim Galore! (https://www.bioinformatics.babraham.ac.uk/projects/trim_galore/) using the –nextera option and a minimum read length of 30 bp. Filtered paired-end reads were aligned to the *M. polymorpha* reference genome (MpTak1_v7.1)[47] using Bowtie2 [48] with the --very-sensitive preset and a maximum fragment length of 2,000 bp. Reads mapping to the chloroplast or mitochondrial genomes were excluded, and only alignments with a mapping quality of ≥30 were retained using SAMtools [49]. PCR duplicates were removed using MarkDuplicates from Picard (https://broadinstitute.github.io/picard/). Accessible chromatin peaks were subsequently called with Genrich in ATAC-seq mode (-j) (https://github.com/jsh58/Genrich).

### Identification of cEEs in tomato and Marchantia

For tomato and Marchantia, exon, coding sequence and gene coordinates were extracted from the corresponding genome annotation files (M82 v1.0 for tomato and MpTak_v7.1 for Marchantia), restricting the analysis to canonical chromosomes (chr1–chr8 in Marchantia and chr1–chr12 in tomato). Exon coordinates included both coding and untranslated regions (UTRs), whereas CDS coordinates were extracted separately to restrict cEE identification to coding sequences. To avoid redundancy arising from alternative transcript isoforms, overlapping exons from transcripts on the same strand were merged using BEDTools merge (-s) to generate a non-redundant exon dataset. To exclude chromatin accessibility associated with promoter-proximal and transcript-end regions, ±100-bp windows were generated around annotated TSSs and transcription end sites (TESs), and exons overlapping these intervals on the same strand were removed using BEDTools intersect (-s -v).

For the permissive analysis, the remaining distal exons were intersected with ATAC-seq peaks using BEDTools intersect, requiring an overlap of at least 1 bp. Among these, only exons that also overlapped an annotated CDS were retained as cEEs. For the stringent analysis, ATAC-seq peak summits, defined as the 1-bp position of maximum accessibility within each peak, were extracted from the narrowPeak files and intersected with the distal exons. Only exons in which the peak summit overlapped an annotated CDS were retained as cEEs. Finally, candidate cEEs were assigned to their respective host genes using BEDTools intersect (-s) and groupby.

### Genome-wide characterization of cEEs

Genomic annotations for Arabidopsis, tomato and Marchantia were retrieved from Ensembl Plants using the txdbmaker/GenomicFeatures packages in R, and CDS and gene coordinates were extracted using the cds() and genes() functions, respectively. cEEs contained within CDS regions were identified using GenomicRanges, and corresponding host genes were assigned based on genomic overlap. Positional duplicates were removed to avoid redundancy arising from alternative transcript isoforms.

For each cEE, its relative position within the host gene was calculated in a strand-aware manner, with 0% and 100% corresponding to the TSSs and TESs, respectively. As the precise sequence corresponding to the cEE within the exon is unknown, the midpoint of the corresponding exon was used as a proxy for its position. cEE occupancy was calculated as the proportion of host-gene length covered by the cEE, and cEEs were classified into four occupancy intervals (0–25%, 25– 50%, 50–75% and 75–100%). The intragenic distribution of cEEs was assessed by kernel density estimation (KDE) for all cEEs and for the subset with ≤25% occupancy, to minimize potential biases arising from large cEEs. Visualizations were generated using ggplot2.

Gene Ontology (GO) enrichment analysis of cEE-associated genes was performed using PlantRegMap [50], testing Biological Process categories against the Arabidopsis genome as background. GO terms with an adjusted *P*-value < 0.05 were considered significantly enriched. Redundancy among enriched GO terms was reduced using REVIGO [51], with the SimRel semantic similarity measure and a medium reduction threshold (0.7). Enriched GO terms were visualized using ggplot2 in R.

Arabidopsis cEEs at selected loci were visualized on the TAIR10 genome using the UCSC Genome Browser and the publicly available EE track hub [16]. For visualization at selected loci, regions containing clusters of predicted TFBSs based on JASPAR motifs or overlapping STARR-seq peaks from published datasets [52] were manually annotated.

### Transcription factor binding-site analyses

Transcription factor binding motifs for Arabidopsis, tomato and Marchantia were obtained from the Plant Transcription Factor Database (PlantTFDB) v5 (https://planttfdb.gao-lab.org/) [53]. Two complementary approaches were used for TFBS analyses. For the identification of individual TFBSs in sequences of interest, a promoter-derived sequence background model was established. Promoter sequences, defined as the 500-bp region upstream and 100-bp downstream of the transcription start site, were retrieved from PlantRegMap[50]. A first-order Markov model was generated from the complete promoter sequence set using fasta-get-markov from the MEME Suite v5.5.9 [54]. Motif occurrences were identified using FIMO [55], scanning both strands with a *P*-value threshold of 1×10^−4^ and using the promoter-derived Markov model to account for background sequence composition. Motif occurrences with a *q*-value <0.05 were retained as significant TFBSs.

For motif enrichment analysis of cEEs, candidate exon sequences were extracted using BEDtools getfasta [56]. Motif enrichment in the sequences of interest was assessed using Simple Enrichment Analysis (SEA; https://meme-suite.org/meme/tools/sea). Shuffled input sequences preserving 3-mer frequencies were used as controls, following SEA default settings. SEA was run using its default E-value threshold (E-value ≤10), after which an additional *q*-value <0.05 threshold was applied to retain significant TFBSs. Retained motifs were grouped by TF family based on PlantTFDB annotations, with representation calculated as the proportion of members represented by at least one retained motif and visualized in R using ggplot2. TF families containing only one member were excluded from the analysis. For Arabidopsis, families were visualized separately according to size (≥15 or <15 members).

### Chromatin interaction, epigenomic and gene-expression analyses

Published chromatin interaction datasets were used to investigate chromatin interactions associated with cEEs in Arabidopsis and tomato. For Arabidopsis, two Micro-C replicates (SRR36568296 and SRR36568295; BioProject PRJNA1392550; under review) were aligned to the TAIR10 reference genome using BWA-MEM and processed with pairtools to parse (mapping quality ≥ 40), sort and deduplicate validate interaction pairs. Replicates were merged, resulting in 289,364,011 read pairs. Multiresolution contact matrices (including 5-kb resolution) were generated with Juicer Tools. Chromatin loops were called at 1-kb resolution using HOMER [57]. For tomato, micro-C data were obtained from GEO (GSE347688) and processed similarly. Aggregated contact analyses were performed on 5-kb resolution matrices using hicAggregateContacts from HiCExplorer [58,59], calculating mean observed/expected intra-chromosomal contacts across 16 bins for cEE-associated genes and comparing them to random genomic control regions. Representative Arabidopsis loops were visualized using the WashU Epigenome Browser (https://epigenomegateway.wustl.edu/browser/).

Gene expression was calculated from published RNA-seq datasets (GEO accessions GSE292906 for Arabidopsis and GSE206365 for tomato) [60,61]. Genes were classified into five groups according to expression level, and the number and/or strength of chromatin loops anchored by cEE-associated genes were compared with those of genes from the different expression groups using Wilcoxon tests. Long-range cEE-associated interactions were defined as interactions spanning >10 kb with a Z-score >1,000 and FDR <0.05, and expression levels of cEE-targeted and non-targeted genes were compared.

For tomato, published ATAC-seq data[61] and histone-modification datasets [62] were additionally analyzed. Signal matrices centered on cEEs (±1 kb) and on the TSSs of cEE host genes (±2 kb) were computed using computeMatrix and visualized as heatmaps and average profiles with deepTools.

### Orthogroup analysis

Orthogroups between Arabidopsis, Marchantia and tomato were inferred using OrthoFinder v2.3.1 [63] with default parameters, using the *A. thaliana* Araport11, *M. polymorpha* Tak1_v7.1 and *Solanum lycopersicum* M82 v1.0 proteomes. For subsequent analyses, orthogroups containing at least one gene from each species were retained. Orthogroups containing genes with cEEs were then identified separately for each species, and their overlap was visualized using the eulerr package in R (https://ceur-ws.org/Vol-2116/paper7.pdf). Statistical significance of the overlap between orthogroups containing cEE-associated genes was assessed using a hypergeometric test, with 12,304 orthogroups defined as the background universe for the tomato– Arabidopsis comparison and 7,086 orthogroups containing at least one gene from each species used as the background universe for the comparison involving Marchantia.

### Phylogenetic analysis

*BRI1* and *BRI1-like* coding sequences from *Arabidopsis thaliana* and *Oryza sativa* were retrieved from Phytozome v14 [64], except those from *Cochlearia groenlandica* [33]. Coding sequences were aligned by codon using ClustalW under default parameters [65]. Phylogenetic relationships were inferred by Maximum Likelihood using the Tamura-Nei nucleotide substitution model [66], with uniform rates among sites, using all 3,936 aligned positions. The initial tree for the heuristic search was selected based on the highest log-likelihood between a Neighbor-Joining tree and the best of 10 random-addition Maximum Parsimony starting trees. Branch support was assessed using the adaptive bootstrap procedure implemented in MEGA v12.1.2 [67], which determined 42 replicates, and support values are shown next to the corresponding branches. The resulting unrooted tree was visualized and edited using iTOL v6 [68].

## Data availability

Scripts used in this project can be found at (https://github.com/rauldenia1/ExonicEnhancers). All relevant information and data of this study are provided in the manuscript and the associated Supplementary Data (including Supplementary Tables 1–20).

## Acknowledgments

We thank Jean-Christophe Mouren and Benoit Ballester (Aix Marseille Univ, INSERM, TAGC, Marseille), and Jaime Pérez-Alemany and Javier Gallego-Bartolomé (IBMCP, Valencia) for insightful early discussions; and Grégory Vert (LRSV, Toulouse) for kindly sharing Gateway plasmids containing different promoters. We also appreciate the assistance of Marisol Gascón at the IBMCP Microscopy Service.

## Author contributions

N.B.T. and C.S.H. conceived and conceptualized the study. N.B.T. performed most of the experiments, with contributions from P.B.-G., A.S.-M., and M.D.-G. R.D.-M. and X.M. carried out all bioinformatic analyses. D.L., M.B., C.S.H., and N.B.T. supervised the study and secured funding. N.B.T. prepared the figures and wrote the manuscript, finalized with C.S.H. and input from all authors. All authors reviewed and approved the final version of the manuscript.

## Funding

This work was supported by grant PID2024-155135NA-I00 from MICIU/AEI/10.13039/501100011033/FEDER, EU, and by the Ramón y Cajal Programme (grant RyC2023-044025-I) from MCIU/AEI/10.13039/501100011033 and FSE+, both awarded to N.B.T.; by institutional grant CEX2025-001626-S (Center of Excellence Severo Ochoa), funded by MCIU/AEI/10.13039/501100011033; and by Swiss National Science Foundation grants 310030_207876 and 3200-0-242995 awarded to C.S.H. R.D.-M. is supported by a predoctoral contract (PREP2024-0025439) associated with grant PID2024-155135NA-I00.

## Competing interests

The authors declare no competing interests.

---

**Suppl. Fig. S1.**
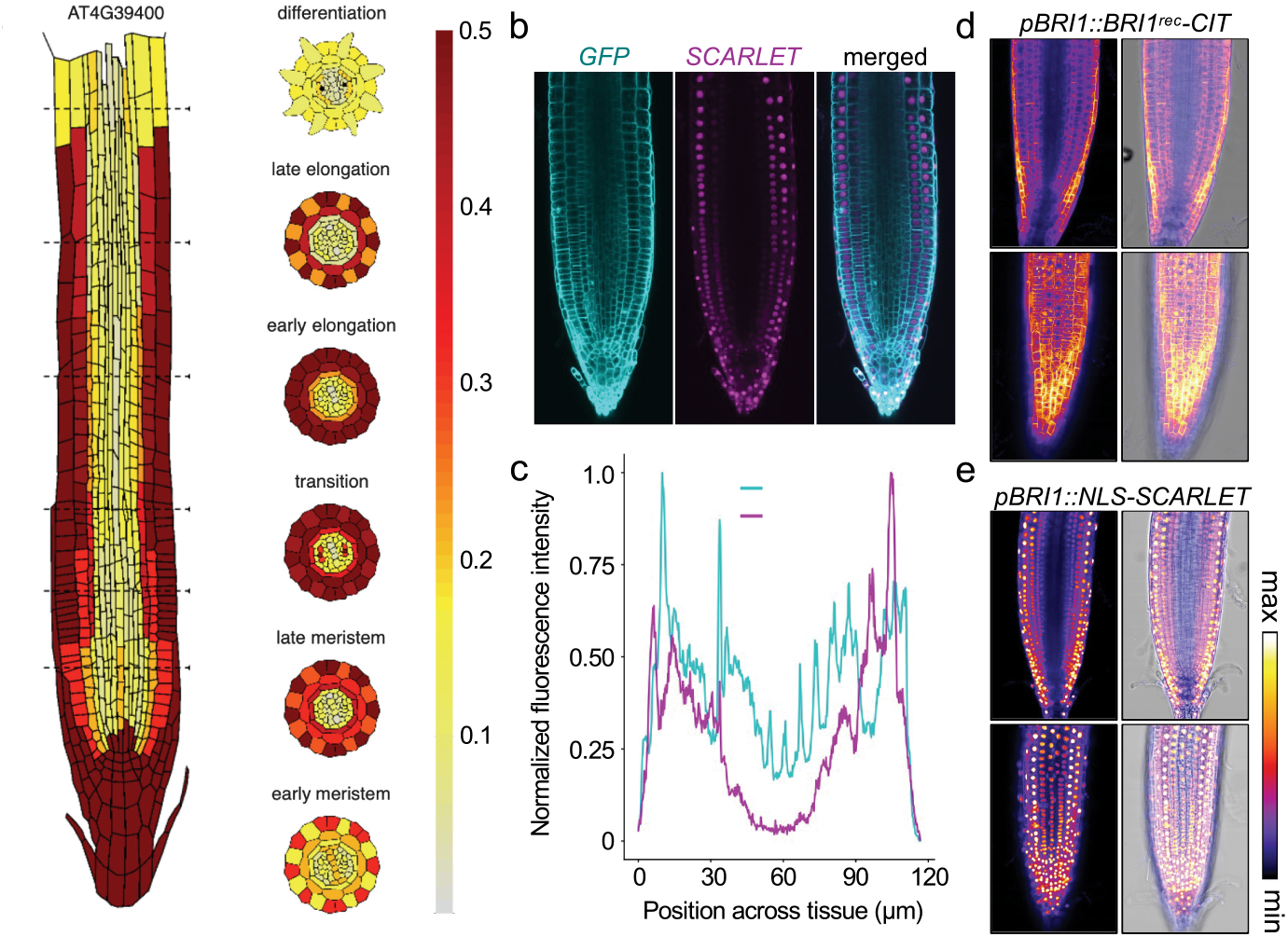
*BRI1* spatial expression extends beyond the promoter-defined domain independently of transgene expression level. **a**, Cell type-specific expression map of endogenous *BRI1* in the Arabidopsis root generated using the Arabidopsis Root Cell Atlas (https://rootcellatlas.org). Longitudinal and transverse sections illustrate the distribution of *BRI1* expression across developmental zones and cell types. The color scale indicates normalized expression levels. **b**, Representative confocal images showing *BRI1*–GFP and *NLS*–SCARLET fluorescence in *pBRI1::BRI1–GFP pBRI1::NLS–SCARLET* seedlings. The white box indicates the approximate region used for fluorescence intensity profiling in c. Scale bar, 20 µm. **c**, Representative fluorescence intensity profiles across the region indicated in b. GFP and NLS–SCARLET fluorescence intensities were independently normalized to their respective minimum and maximum values. **d**, Confocal images of representative *pBRI1::BRI1*^*rec*^*–CITRINE* transformants exhibiting higher fluorescence intensity than those shown in Fig. 1c. Upper panels show optical sections through the quiescent center (QC), and lower panels through the epidermis. Fluorescence intensity is shown in false color (left) and merged with transmitted-light images (right). Cell layers are indicated (ep, epidermis; c, cortex; en, endodermis; p, pericycle). Scale bars, 20 µm. **e**, Confocal images of representative *pBRI1::NLS–SCARLET* reporter lines exhibiting higher fluorescence intensity than the line shown in Fig. 1e. Images are displayed as in d. Scale bars, 20 µm.

**Suppl. Fig. S2.**
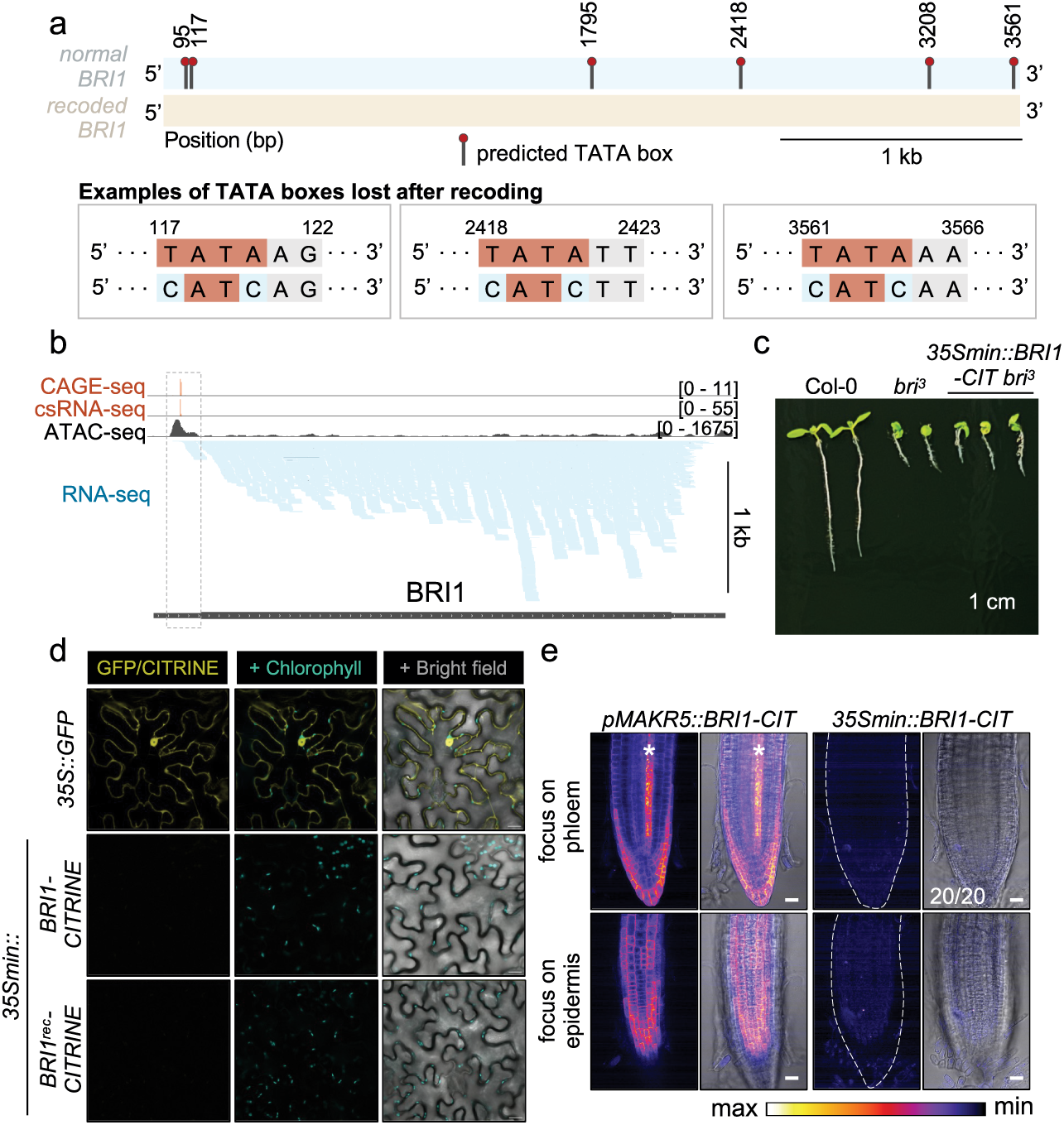
The regulatory activity of the *BRI1* coding sequence is not explained by cryptic promoter activity or alternative transcription initiation. **a**, Schematic comparison of the native and synonymously recoded *BRI1* coding sequences showing predicted TATA box motifs. Representative TATA box motifs lost upon synonymous recoding are shown below. **b**, Genome browser view of the *BRI1* locus showing CAGE-seq, csRNA-seq, ATAC-seq and RNA-seq datasets. No evidence of transcription initiation was detected within the *BRI1* coding sequence. **c**, Representative 6-day-old seedlings of Col-0, *bri*^*3*^, and *35Smin::BRI1–CIT bri*^*3*^. Expression of the *BRI1* coding sequence downstream of a minimal *35S* promoter failed to complement the *bri*^*3*^ phenotype. Scale bar, 1 cm. **d**, Transient expression of *35Smin::BRI1–CITRINE* and *35Smin::BRI1*^*rec*^*–CITRINE* in *Nicotiana benthamiana* leaves. CITRINE fluorescence was undetectable from both *BRI1* constructs, whereas *35S::GFP*, infiltrated in parallel as an infiltration control, produced robust fluorescence. CITRINE fluorescence, chlorophyll autofluorescence and merged fluorescence/transmitted-light images are shown. More than 10 leaf discs were examined per construct. **e**, Confocal images of representative *pMAKR5::BRI1–CIT* and *35Smin::BRI1–CITRINE* Arabidopsis roots. Upper panels show optical sections through the phloem and lower panels through the epidermis. Fluorescence intensity is shown in false color (left) and merged with transmitted-light images (right). The number of independent transformants analyzed is indicated. Asterisks denote the phloem. Dashed lines outline the root. Scale bars, 20 µm.

**Suppl. Fig. S3.**
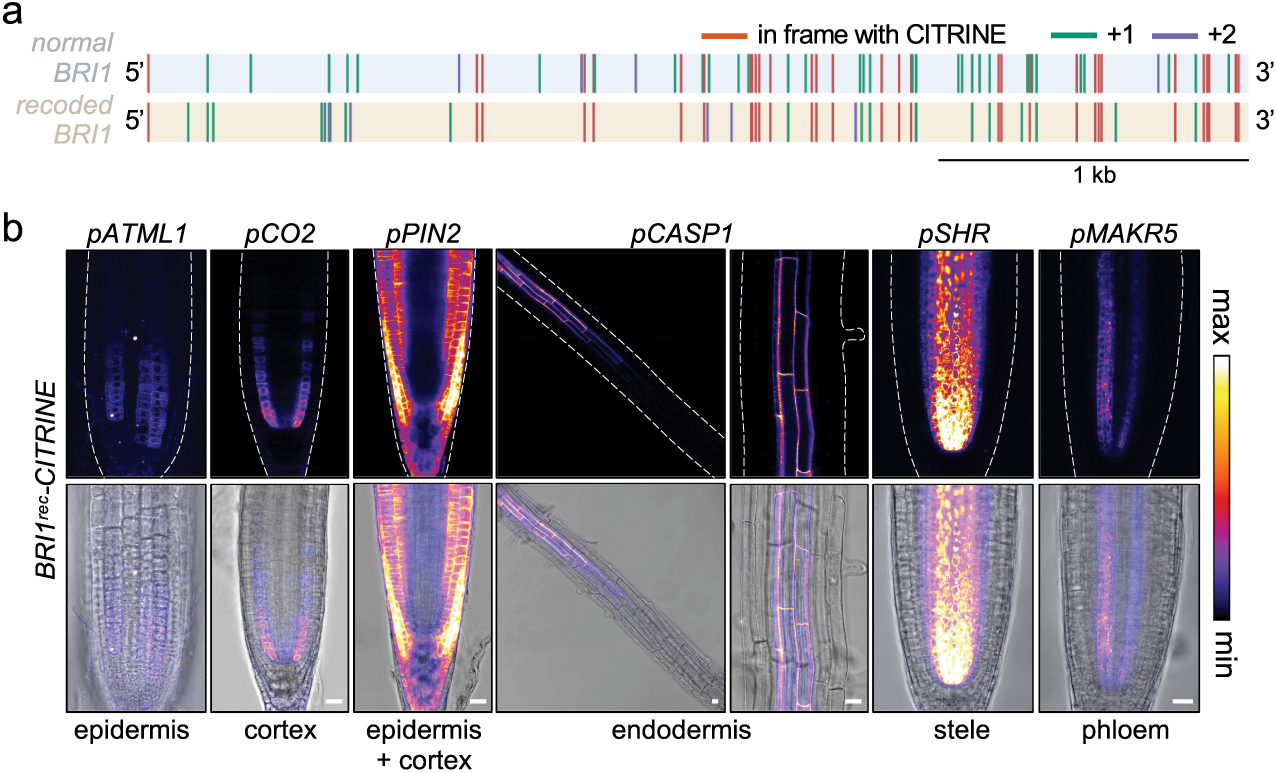
Synonymous recoding does not alter translation initiation and permits BRI1 production across multiple root cell types. **a**, Schematic representation of the positions of ATG codons in the three reading frames relative to the *BRI1*–CITRINE fusion in the native and synonymously recoded *BRI1* coding sequences. Synonymous recoding neither altered the canonical translational start site nor introduced additional in-frame ATG codons that could produce alternative BRI1–CITRINE fusion proteins. **b**, Representative confocal images of Arabidopsis roots expressing *BRI1*^*rec*^–CIT under the control of the epidermis-specific *pATML1*, cortex-specific *pCO2*, epidermis- and cortex-specific *pPIN2*, endodermis-specific *pCASP1*, stele-specific *pSHR*, and phloem-specific *pMAKR5* promoters. Upper panels show fluorescence intensity in false color and lower panels show merged fluorescence and transmitted-light images. *BRI1*^*rec*^–CIT accumulation was observed in all examined cell types, indicating that synonymous recoding does not impair protein production. Scale bars, 20 µm.

**Suppl. Fig. S4.**
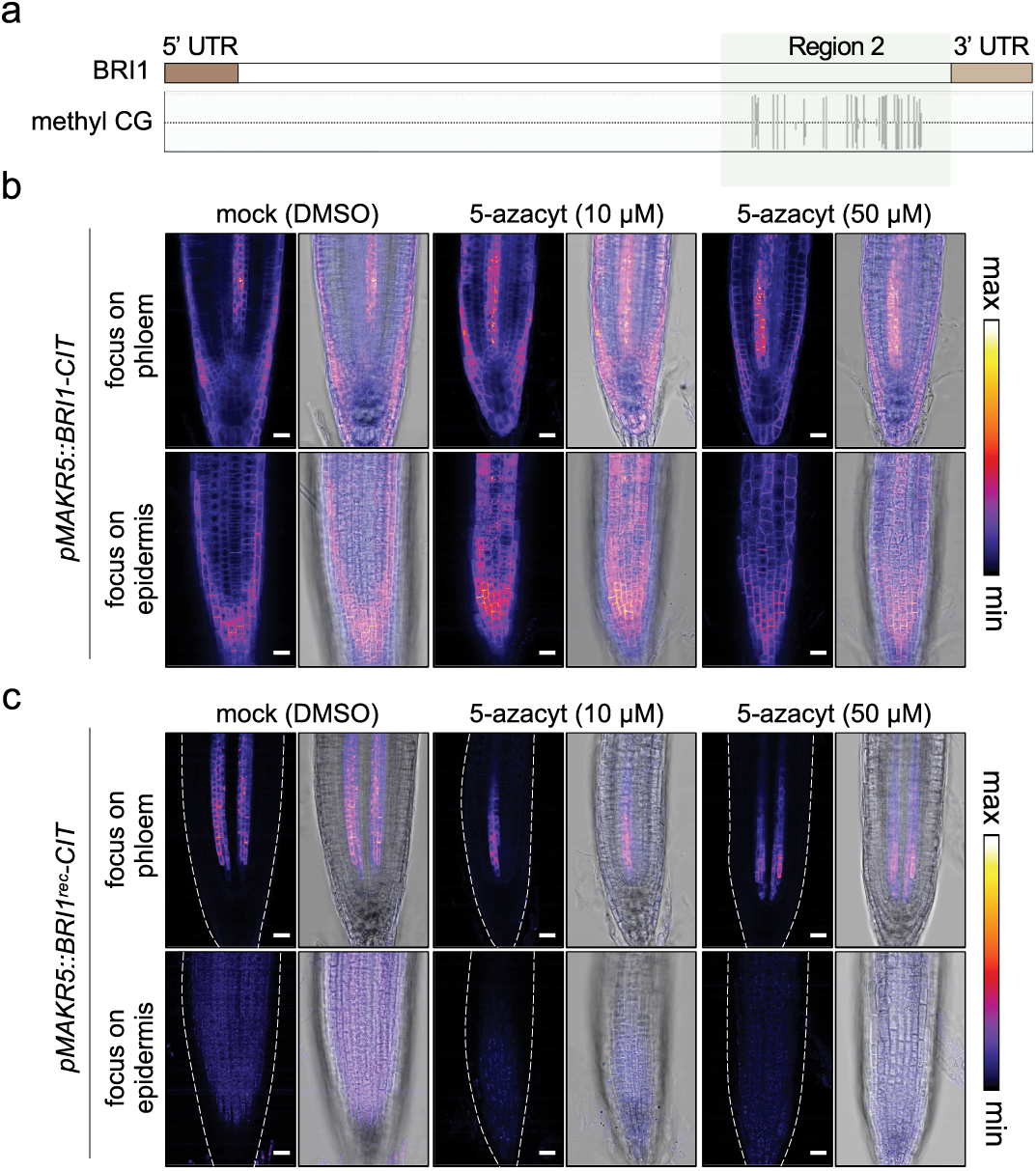
DNA methylation is not required for the coding sequence-dependent regulatory activity of *BRI1*. **a**, Schematic representation of CG DNA methylation in the *BRI1* locus (salk.edu). Region 2, encoding the intracellular portion of BRI1 analyzed in Fig. 2, is highlighted. **b**, Confocal images of representative *pMAKR5::BRI1–CIT* roots treated with mock (DMSO) or 5-azacytidine (10 or 50 µM). Upper panels show optical sections through the phloem and lower panels through the epidermis. Fluorescence intensity is shown in false color (left) and merged with transmitted-light images (right). Scale bars, 20 µm. **c**, Confocal images of representative *pMAKR5::BRI1*^*rec*^*–CIT* roots treated as in b. Images are displayed as in b. In both native and recoded *BRI1* lines, 5-azacytidine treatment resulted in shorter root meristems. Scale bars, 20 µm.

**Suppl. Fig. S5.**
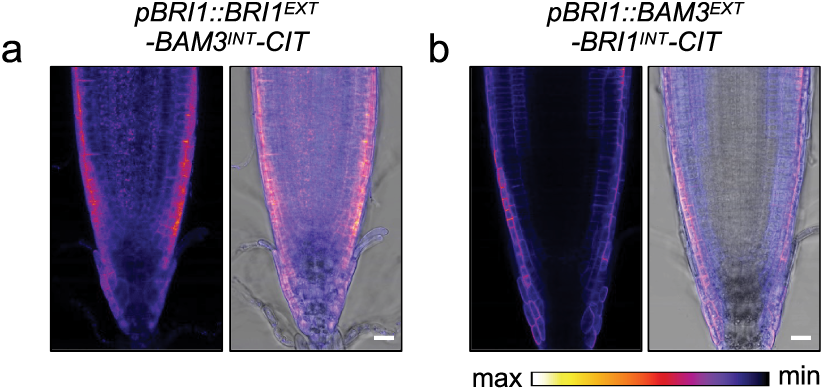
The *BRI1* coding sequence-embedded regulatory activity is transferable to a heterologous coding sequence context. **a**, Confocal images of Arabidopsis roots expressing *pBRI1::BRI1*^*EXT*^*–BAM3*^*INT*^*–CIT*. Fluorescence intensity is shown in false color (left) and merged with transmitted-light images (right). Scale bars, 20 µm. **b**, Confocal images of Arabidopsis roots expressing *pBRI1::BAM3*^*EXT*^*–BRI1*^*INT*^*–CIT*. Images are displayed as in a. Scale bars, 20 µm.

**Suppl. Fig. S6.**
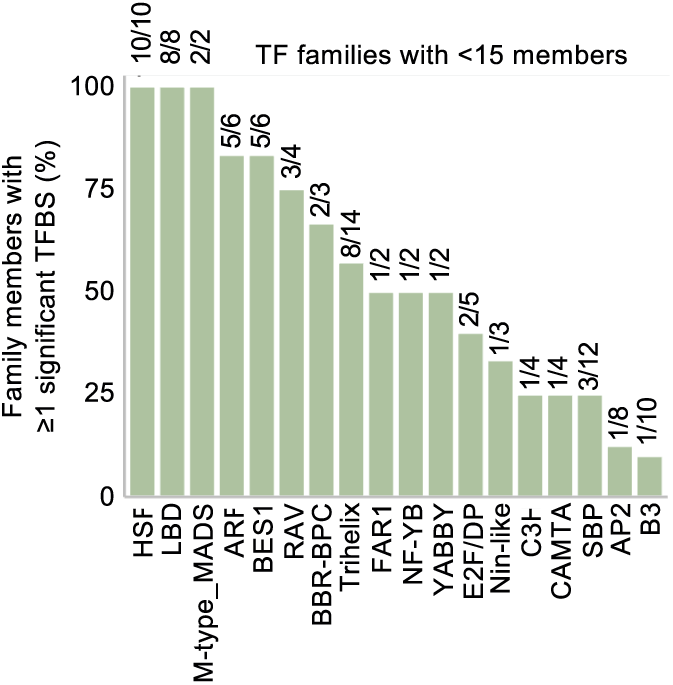
Representation of transcription factor families with fewer than 15 members in Arabidopsis candidate exonic enhancers. Relative representation of TF families comprising fewer than 15 members among candidate exonic enhancers. Bar height represents the percentage of TFs within each family represented by at least one significant TFBS, with numbers indicating the corresponding number of TFs relative to the total number of TFs in that family.

**Suppl. Fig. S7.**
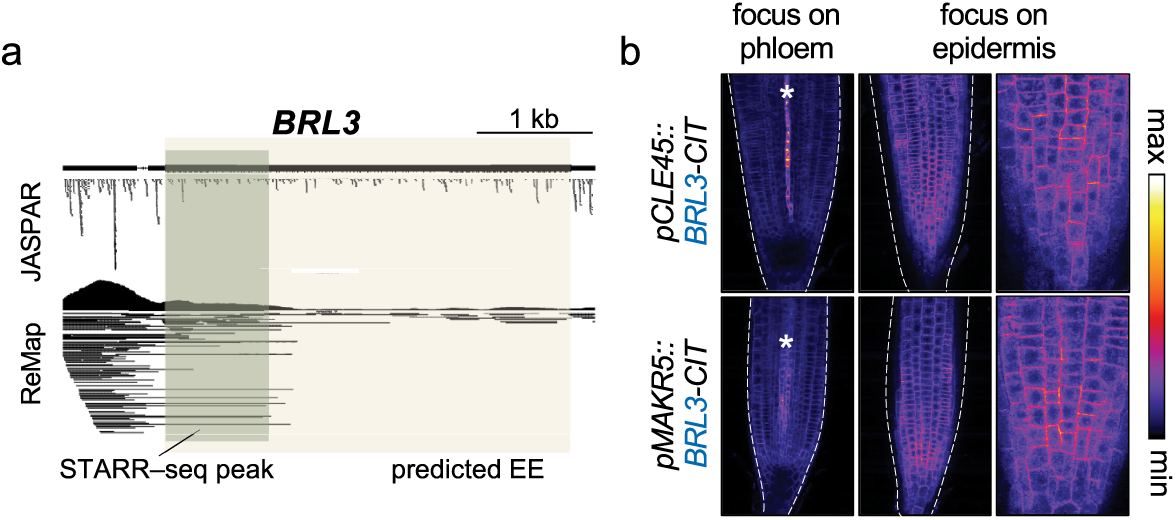
Coding sequence-dependent spatial regulation extends to the brassinosteroid receptor BRL3. **a**, Genome browser view of the *BRL3* locus showing predicted transcription factor binding sites (JASPAR), ReMap transcription factor binding data, and a STARR-seq peak overlapping the predicted EE. **b**, Confocal images of Arabidopsis roots expressing *pCLE45::BRL3–CITRINE* or *pMAKR5::BRL3–CITRINE*. Left panels show optical sections through the phloem and middle panels through the epidermis. Right panels show higher-magnification views of the epidermis. Fluorescence intensity is shown in false color. Asterisks denote the phloem. Dashed lines outline the root. Scale bars, 20 µm.

**Suppl. Fig. S8.**
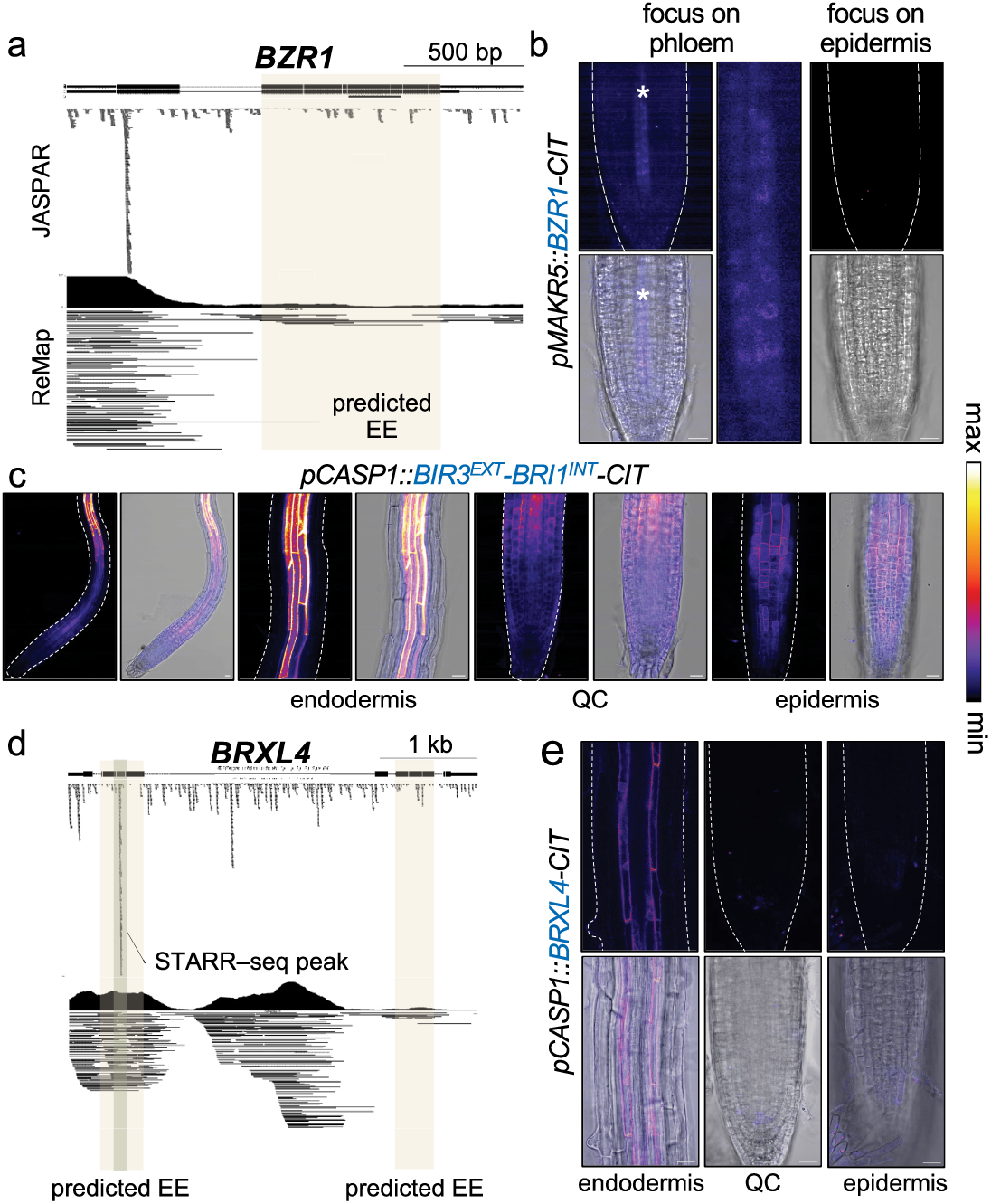
Additional predicted exonic enhancers do not contribute to expanded spatial expression patterns. **a**, Genome browser view of the *BZR1* locus showing predicted transcription factor binding sites (JASPAR) and ReMap transcription factor binding data. The predicted EE is highlighted. **b**, Confocal images of Arabidopsis root apices expressing *pMAKR5::BZR1–CITRINE*. Left panels show optical sections through the phloem and right panels through the epidermis. The middle panel shows a higher-magnification view of the phloem. Fluorescence intensity is shown in false color (upper panels) and merged with transmitted-light images (lower panels). Asterisks denote the phloem. Dashed lines outline the root. Scale bars, 20 µm. In contrast to *BRI1* and *BRL* receptors, *BZR1* expression remains confined to the promoter-defined domain. **c**, Confocal images of Arabidopsis root apices expressing *pCASP1::BIR3*^*EXT*^*–BRI1*^*INT*^*–CITRINE*, included as a positive control for coding sequence-dependent expansion beyond the *pCASP1*-defined expression domain. Note that expression extends beyond the *CASP1* promoter-defined endodermal domain into the root meristem. QC, quiescent center. Fluorescence intensity is shown in false color and merged with transmitted-light images. Dashed lines outline the root. Scale bars, 20 µm. **d**, Genome browser view of the *BRXL4* locus showing predicted transcription factor binding sites (JASPAR), ReMap transcription factor binding data and a STARR-seq peak overlapping a predicted exonic enhancer. Predicted exonic enhancers are indicated. **e**, Confocal images of Arabidopsis root apices expressing *pCASP1::BRXL4–CITRINE*. Left, optical sections through the endodermis. Middle, optical sections through the quiescent center (QC). Right, optical sections through the epidermis. Fluorescence intensity is shown in false color (upper panels) and merged with transmitted-light images (lower panels). Dashed lines outline the root. Scale bars, 20 µm.

**Suppl. Fig. S9.**
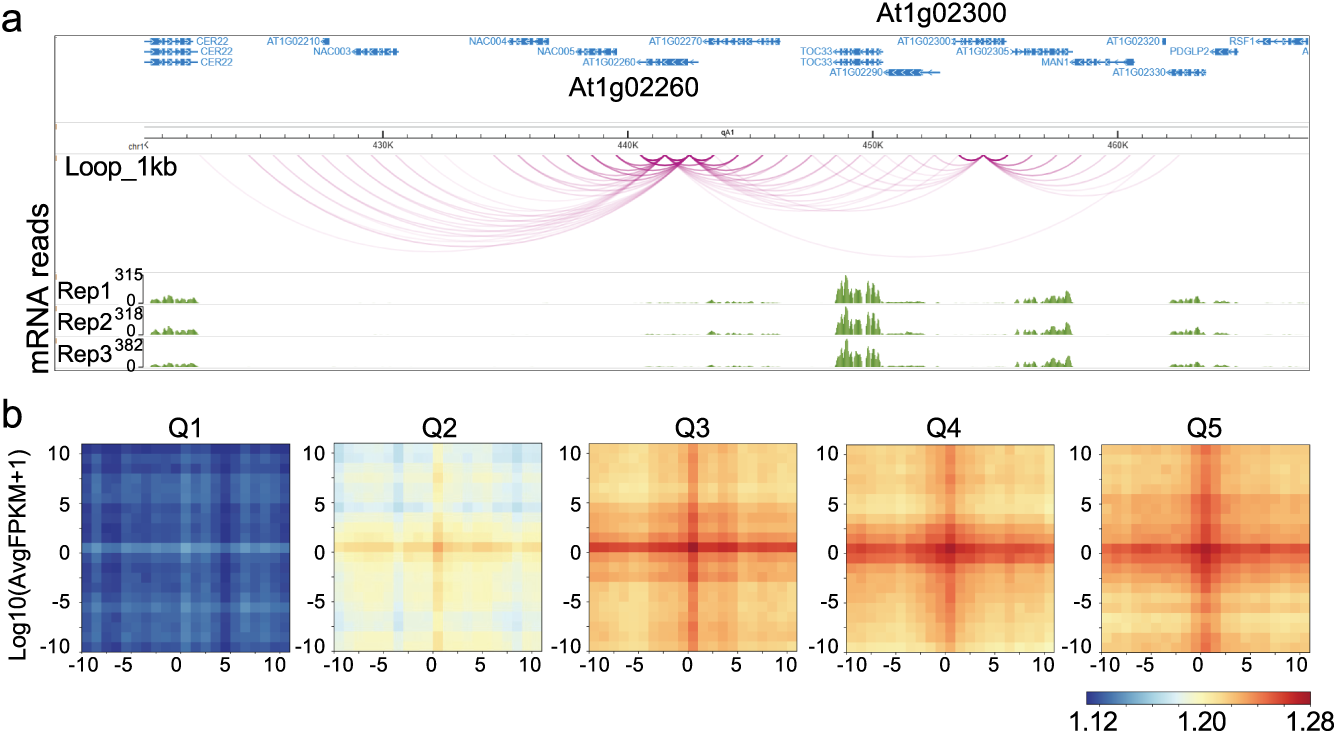
Chromatin interaction profiles associated with candidate exonic enhancer-containing genes. **a**, Representative genomic view of Micro-C chromatin interactions at a genomic region containing two candidate exonic enhancer (cEE)-containing genes, At1g02260 and At1g02300. Chromatin loops identified at 1-kb resolution are shown together with gene annotations and RNA-seq coverage across three biological replicates. **b**, Aggregate contact maps of Arabidopsis genes grouped into five expression quintiles (Q1–Q5) based on RNA-seq expression levels, from lowest (Q1) to highest (Q5) expression.

**Suppl. Fig. S10.**
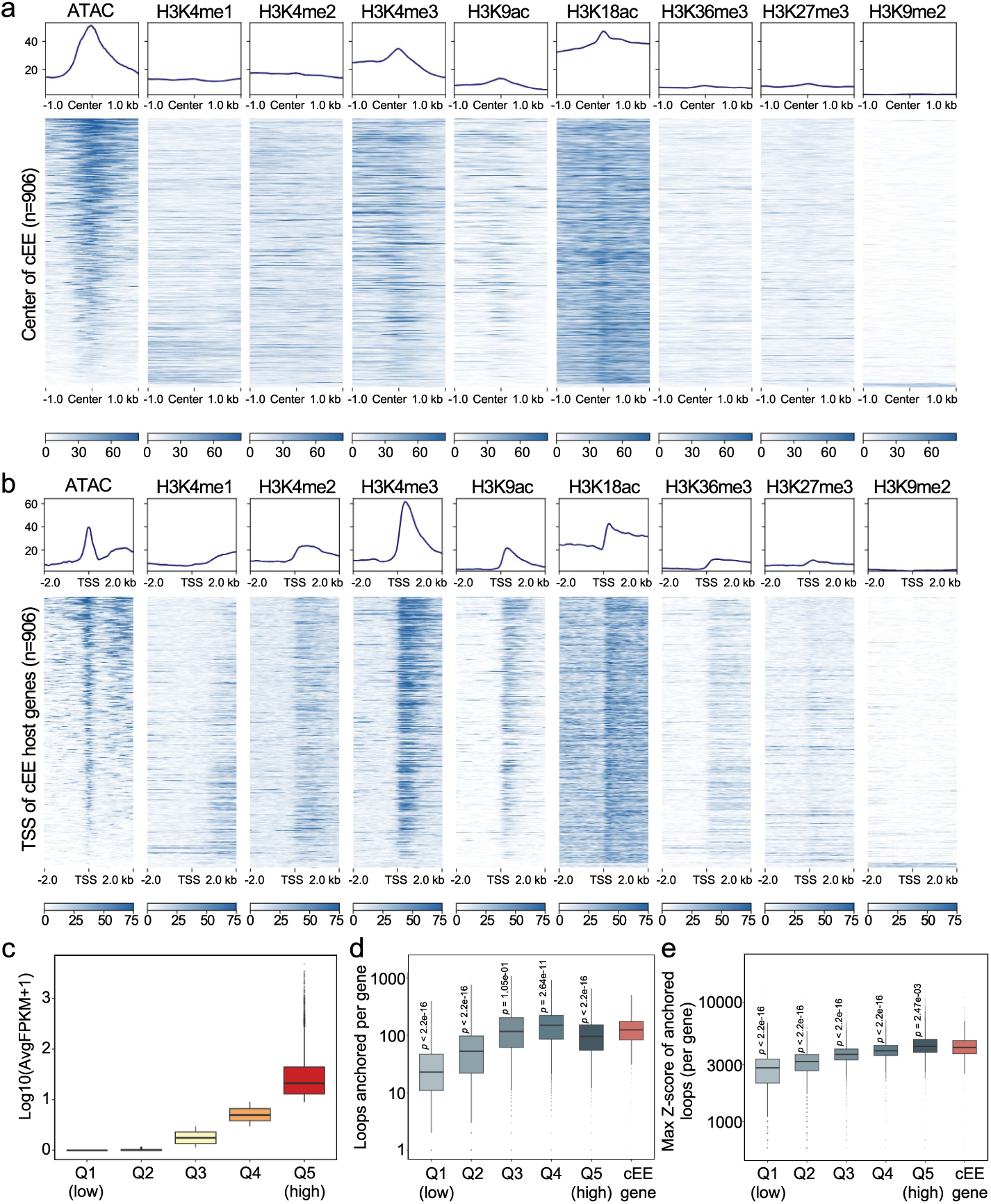
Epigenomic and chromatin interaction features associated with candidate exonic enhancer-containing genes in tomato. **a**, Signal profiles and heatmaps of ATAC-seq and the indicated histone modifications centered on candidate exonic enhancers (cEEs) in tomato (*Solanum lycopersicum*). Each heatmap spans ±1 kb from the cEE center. **b**, Signal profiles and heatmaps of ATAC-seq and the indicated histone modifications around the transcription start sites (TSSs) of cEE-containing genes. Each heatmap spans ±2 kb from the TSS. **c**, Distribution of gene expression levels used to define the five expression quintiles (Q1–Q5) for subsequent chromatin interaction analyses. **d**, Number of chromatin interactions anchored at cEE-containing genes and genes grouped by expression quintile (Q1–Q5). **e**, Maximum *Z*-score of anchored chromatin loops per gene for cEE-containing genes and genes grouped by expression quintile (Q1–Q5). For d and e, *p* values were determined using Wilcoxon rank-sum tests comparing cEE-containing genes with each expression quintile.

## Notes

### Competing Interest Statement

The authors have declared no competing interest.

